# Single cell multi-omics analysis of chromothriptic medulloblastoma highlights genomic and transcriptomic consequences of genome instability

**DOI:** 10.1101/2021.06.25.449944

**Authors:** R. Gonzalo Parra, Moritz J Przybilla, Milena Simovic, Hana Susak, Manasi Ratnaparkhe, John KL Wong, Verena Körber, Philipp Mallm, Martin Sill, Thorsten Kolb, Rithu Kumar, Nicola Casiraghi, David R Norali Ghasemi, Kendra Korinna Maaß, Kristian W Pajtler, Anna Jauch, Andrey Korshunov, Thomas Höfer, Marc Zapatka, Stefan M Pfister, Oliver Stegle, Aurélie Ernst

**Affiliations:** European Molecular Biology Laboratory, Genome Biology Unit, Heidelberg, Germany.; Division of Computational Genomics and Systems Genetics, German Cancer Research Center (DKFZ), Heidelberg, Germany; Wellcome Sanger Institute, Wellcome Trust Genome Campus, Cambridge, UK; Group “Genome Instability in Tumors”, German Cancer Research Center (DKFZ), Heidelberg, Germany; Faculty of Biosciences, Heidelberg University; Division of Molecular Genetics, German Cancer Research Center (DKFZ), Heidelberg, Germany; Division of Theoretical Systems Biology, German Cancer Research Center (DKFZ), Heidelberg, Germany; Single-cell Open Lab, German Cancer Research Center (DKFZ) and Bioquant, Heidelberg, Germany; Hopp Children’s Cancer Center (KiTZ), Heidelberg, Germany. Pediatric Neurooncology, German Cancer Consortium (DKTK) and German Cancer Research Center (DKFZ), Heidelberg, Germany.; Department of Pediatric Hematology and Oncology, University Hospital, Heidelberg, Germany; Institute of Human Genetics, University of Heidelberg, Heidelberg, Germany; Clinical Cooperation Unit Neuropathology, DKFZ, Department of Neuropathology, Heidelberg University Hospital

## Abstract

Chromothripsis is a form of genome instability, whereby a presumably single catastrophic event generates extensive genomic rearrangements of one or few chromosome(s). However, little is known about the heterogeneity of chromothripsis across different clones from the same tumor, as well as changes in response to treatment. We analyzed single-cell genomic and transcriptomic alterations linked with chromothripsis in human p53-deficient medulloblastoma (n=7). We reconstructed the order of somatic events, identified early alterations likely linked to chromothripsis and depicted the contribution of chromothripsis to malignancy. We characterized subclonal variation of chromothripsis and its effects on double-minute chromosomes, cancer drivers and putatively druggable targets. Furthermore, we highlighted the causative role and the fitness consequences of specific rearrangements in neural progenitors.

**Graphical Abstract:** 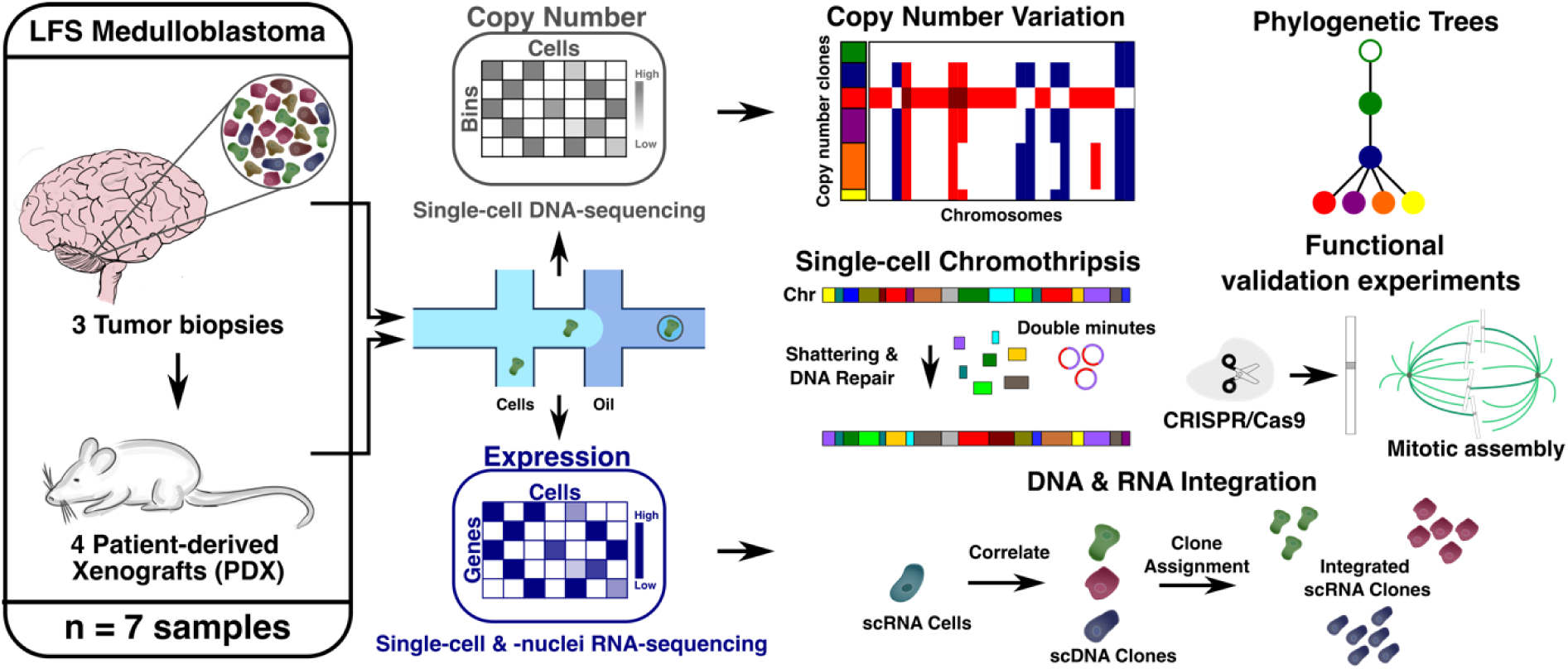

## INTRODUCTION

Chromothripsis is a type of genome instability, by which a presumably single catastrophic event leads to substantial genomic rearrangements of one or a few chromosome(s) ^1,2^. Generally considered as an early event in the evolution of a tumor, chromothripsis likely plays a causative role in the development of a number of tumors by generating multiple genomic aberrations simultaneously. In line with this, rearrangements due to chromothripsis were detected in more than 25% of cancer patients in two large pan-cancer studies ^3,4^. In specific tumor types or molecular subgroups, the prevalence for chromothripsis reaches 100%, such as in medulloblastoma with germline *TP53* mutations (Li-Fraumeni syndrome, LFS), which is the focus of this study. Chromothripsis is linked with poor prognosis for these patients, as in a number of other tumor types ^2,5–7^. In the context of *TP53* mutations in the germline, it is conceivable that multiple chromothriptic (CT) events may occur in different cells and most of them are not selected for and therefore undetected.

Longitudinal studies on the evolution of CT chromosomes between matched primary and relapsed tumors showed that CT patterns may be either i) stabilized ii) eliminated or iii) undetected at initial diagnosis but present in the relapsed tumor ^4,8,9^. Importantly, elimination as well as newly detected CT chromosomes suggest that a subset of tumor cells in the initial tumor may potentially lack or already carry the CT chromosome. These findings question the paradigm that chromothripsis is a single early event in tumor development, which would imply that the CT chromosome would be present in the vast majority of the tumor cells. However, conclusive indications for the same CT patterns across cells within a tumor and for the fraction of cells carrying a given CT chromosome are rare, as rearrangements due to chromothripsis have not been analyzed at single-cell resolution in tumor tissue to date. As chromothripsis was shown to drive tumor development through the activation of oncogenes and the disruption of tumor suppressor genes ^4,10^, it is essential to assess the extent of clonal heterogeneity affecting drivers and therapeutic targets within CT tumors. In addition, chromothripsis was linked to compromised function of essential factors such as p53, ATM and critical DNA repair proteins, suggesting that the inactivation of specific pathways and checkpoints may facilitate the occurrence of CT events and/or the survival of the cells after such an event ^2,11,12^. However, direct evidence of potential enabling mechanisms is limited.

Sequencing of cultured cells showed that processes such as mitotic errors, micronuclei formation, centromere inactivation, chromatin bridges but also telomere dysfunction can cause a range of rearrangements, including chromothripsis ^13–17^. Although modelling chromothripsis in cell culture systems has allowed putative mechanisms to be proposed, the way in which spontaneous CT events occur in human cells remains largely unknown. It is unclear to which extent mechanisms derived from artificially inducing chromothripsis *in vitro* reflect CT events in human cancer. Single-cell DNA sequencing studies in the context of chromothripsis are only beginning to emerge. Pellman and colleagues reported mechanistic insights into the generation of complex rearrangements from sequencing cultured clones and single cells from the retinal pigment epithelial (RPE1) cell line ^16,17^. Korbel and colleagues investigated structural variation in the RPE1 cell line and showed complex rearrangements for one leukemia sample as a proof-of-principle (approximately 80 cells) ^18^. To study this process *in vivo*, we set off to perform the first characterization of the heterogeneity in CT patterns across tumors.

Here, we leverage bulk and single-cell sequencing assays, combined with fluorescence in situ hybridization (FISH), immunofluorescence analyses and CRISPR/Cas9 knockouts to investigate the origins and functional consequences of chromothripsis in LFS medulloblastoma. We generated shallow single-cell DNA- and single-cell RNA-seq profiles from 757 and 22,500 cells from 7 LFS medulloblastoma samples, respectively. For the first time, we demonstrate the ability to detect CT events at the single-cell level in tumors, further unraveling the extent of intra-tumor heterogeneity with clonal resolution. In addition, we highlight potential mechanisms for the formation of double-minute chromosomes. Using our single-cell RNA-seq (scRNA-seq) data, we characterize the cell type landscape and investigate differences to non-CT medulloblastoma. By integrating single-cell DNA-seq (scDNA-seq) and scRNA-seq information based on somatic copy number profiles estimated from both data modalities, we shed light into potential transcriptomic consequences of chromothripsis and its impact on tumor evolution. Finally, we identified a putative role for the SETD2 methyltransferase in the early stages of the development of CT medulloblastomas using functional analyses in neural stem cells.

## RESULTS

To determine how chromothripsis contributes to inter-cell genetic heterogeneity, generating oncogenic drivers that increase cell fitness and hence tumor aggressiveness, we performed single-cell DNA and RNA sequencing of brain tumors with chromothripsis (n=7, including patient tumors and patient-derived xenograft (PDX) models, primary and relapsed, see graphical abstract & **Supplementary Table 1**). We combined the resolution from single-cell genome and transcriptome analyses with a population-scale bulk deep sequencing cohort (n=227), to generate a map of the landscape of alterations linked with chromothripsis. We focused on medulloblastoma in patients with germline *TP53* mutations (LFS), a childhood brain tumor with extremely poor prognosis. As chromothripsis is present in close to 100% of these cancers ^2,19^, medulloblastomas in LFS patients constitute a paradigm for the understanding of this phenomenon.

### Rearrangements due to chromothripsis can be detected in single tumor cells

With chromothripsis being present in all LFS medulloblastomas analyzed so far ^2,19^, we asked whether we could characterize the underlying subclonal heterogeneity using single-cell sequencing. Initially, we considered the primary tumor LFS_MBP (**Fig. 1a-e**, **Supplementary Fig. 1 a-d**) from which the largest numbers of cells were captured by single-cell DNA-seq (scDNA-seq) analysis (N=404 cells, 10x Genomics Chromium Single Cell CNV, **Supplementary Table 1,** see **Methods** for Quality Control criteria). The resolution of the scDNA-seq data was adequate to identify CT chromosomes (107 to 682 reads/Mbp per cell, median 427, **Supplementary Table 1)**, with aggregate profiles (pseudo-bulk) showing high concordance with bulk somatic copy number profiles derived from whole-genome sequencing (WGS) data (**Fig. 1a** - LFS_MBP-Nuclei and **Supplementary Fig. 2a** - additional samples). Rare CNVs present in the germline (e.g. gain on chromosome 1) were detected in CNV profiles derived from scDNA-seq but not visible in bulk somatic copy number profiles (the latter adjusted for CNVs present in matched normal control blood) (**Fig. 1a**). To study clonal heterogeneity, we reduced noise by adjusting for rare aberrant ploidy estimates at individual loci in the single cell CNV dataset (**Supplementary Fig. 1e, Methods**), followed by hierarchical clustering of the resulting single-cell CNV profiles (N=2 to 6 clones per tumor, **Supplementary Fig. 2a-e**). Interestingly, this analysis identified mirrored copy-number patterns in pairs of clones, possibly originating from one mitotic error (e.g. **Fig. 1a**, chromosome 7 in clones C4 and C5, highlighted in blue - LFS_MBP-Nuclei; **Supplementary Fig. 2b** - LFS_MBP-PDX). Such mirrored copy-number patterns with regions gained in one clone but lost in the mirrored clone remain undetected by bulk sequencing, which quantifies average copy-number profiles. Leveraging the aggregate information across all cells assigned to a clone, we estimated phylogenetic clonal trees based on the distances of the CNV profiles between individual clones (**Fig. 1b**; **Supplementary Fig. 2a-f,** see **Methods**). Comparison of the underlying CNV profiles between clones identified clone-specific cancer drivers (e.g. focal gain of *MYCN,* **Fig. 1a** - LFS_MBP-Nuclei**, Supplementary Fig. 2b** - LFS_MBP-PDX). When extending this analysis to all 7 samples, we observed marked variation between samples, including between matched primary and relapsed tumor profiles from the same patient, as well as between patient tumors and matched patient-derived xenograft (PDX) models (**Fig. 1c**). For instance, only two out of six clones present in LFS_MBP (primary patient tumor) displayed corresponding clones in the derived PDX sample (LFS_MBP_PDX) and none of the six major clones identified in LFS_MBP displayed a closely related clone in the matched relapse (LFS_MB1R, relapsed patient tumor) (**Fig. 1c**, Methods). Altogether, this indicates substantial intra and inter genetic heterogeneity in CT medulloblastomas.

**Figure 1.**
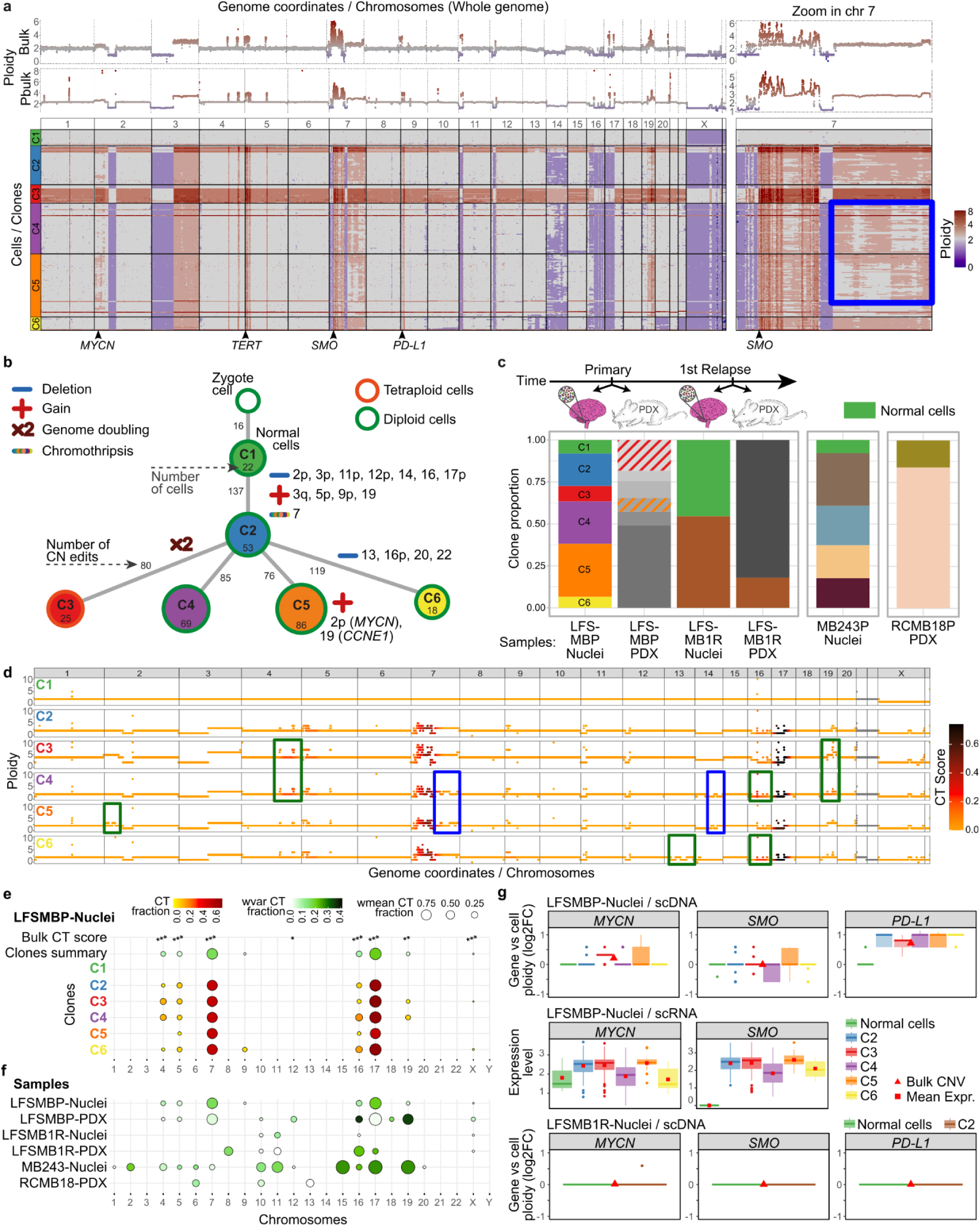
Genetic heterogeneity in medulloblastomas with chromothripsis. **a,** Overview of copy number variation and clonal substructure estimated using scDNA-seq (LFS_MBP, primary tumor, nuclei). From top to bottom: somatic copy number profiles from bulk DNA-seq (Bulk), aggregate scDNA-seq (Pbulk), and CNV profiles of individual cells grouped into 6 major clones based on CNV clustering. Zoom-in view displays a major genomic rearrangement on chromosome 7, indicating concordance between aggregate single-cell and bulk profiling, but also the presence of compensatory CNV changes that can be exclusively detected using single-cell sequencing (blue box). **b,** Estimated phylogenetic clonal tree for LFS_MBP-primary (evolutionary edit distance; Methods). Top node denotes a synthetic zygote from which clones are laid out according to their respective CNV edit distance (evaluated in 20 kb genomic intervals). Numbers on lines denote the number of CNV changes; numbers in nodes denote the number of cells assigned to each clone; selected events are highlighted. **c,** Clonal composition across all 7 samples, displaying clonal composition as in **b**. Bar height denotes the relative proportion of cells assigned to a clone. Matched clones with concordant CNV profiles across samples from the same patient are annotated with matching color; suggestive correspondences with striped patterns (see Methods); normal cell populations are shown in green. **d,** CT scoring for individual clones as in **c** for LFS_MBP-primary, across genomic coordinates. Bin CT fraction denotes the fraction in which a 20kb segment is scored as chromothriptic (50MB window for CT assessment; 20 kb step size; requiring 10 switches; Methods). Selected clone-specific CNVs and CT events are highlighted in green, mirror-like patterns in blue. Chromosomes 21 and 22 are omitted due to chromosome size (grey). The focal gain on chromosome 1 also detected in normal cells is present in the germline. The focal gain on chromosome 16 also detected in normal cells is close to the centromere. **e,** Analysis of CT heterogeneity in LFS_MBP-primary with chromosomal resolution. From top to bottom: Chromosomes with CT events detected in bulk WGS (based on ShatterSeek; Methods, * p =1.0e-07, ** p = 1.07E-07, *** p =2.22E-16), aggregated CT fractions (fraction of 20kb segments with evidence for CT in a given chromosome; 50MB window for CT assessment; 20 kb step size; Methods), Clones summary aggregate statistics and chromosomal CT fractions for individual clones. Aggregate statistics node size denotes the average CT fraction across clones; color denotes the CT fraction variance across clones. High variance indicates inter clonal CT heterogeneity. **f,** Clone summary statistics of CT as in **e** for all 7 samples, displaying a high degree of variability in the CT signatures across samples. **g,** Analysis of putatively druggable targets for LFS_MB, focusing on representative target genes. From top to bottom: log fold change between the CNV profile for a given gene locus and the median CNV values of this chromosome (focal gains) for each clone (LFSMBP-Nuclei), estimated expression level per clone for expressed genes based on scRNA-seq integration (see Methods and Fig. 4a); log fold change between the CNV profile for a given gene locus and the median CNV values of this chromosome (focal gains) for each clone (LFSMB1R-Nuclei). Bulk CNV values shown as red triangles. The data are represented as boxplots where the middle line corresponds to the median, the lower and upper edges of the box correspond to the first and third quartiles, the whiskers represent the interquartile range (IQR) ×1.5 and beyond the whiskers are outlier points.

Next, we adapted chromothripsis scoring strategies designed for bulk sequencing that estimate the fraction of chromothripsis per chromosome ^3,4^ to single-cell profiles, considering CNV profiles aggregated at the level of clones (**Fig. 1d-f, Supplementary Table 2,** see Methods). In addition to clonal CT events in each sample (e.g. chromosomes 7 and 17 in LFS_MBP-Nuclei, **Fig. 1d-e**), this revealed heterogeneity across clones for some of the CT chromosomes, which was robustly identified also on subsampled data (**Supplementary Fig. 2f, Methods**). In essence, this indicates subclonal heterogeneity in chromothripsis, which contradicts the common view of chromothripsis as a single early event. Furthermore, we confirmed selected subclonal CT events by targeted reanalysis of available bulk WGS data from the same sample and comparisons between primary tumors and derived samples (**Supplementary Fig. 3)**. For example, the subclonal chromothripsis event on chromosome 8 in sample LFS_MB1R, barely detectable in the bulk WGS, was only present in a subset of cells in the single-cell data. However, in the matched PDX model, this minor clone with chromothripsis on chromosome 8 became dominant, likely due to selection in the xenograft (**Supplementary Fig. 3)**.

Subsets of CNVs are non-randomly associated with lineage expansion and potentially have functional impact, also in terms of treatment response and resistance. In line with this, we identified focal gains and high-level amplifications affecting putative drug targets, and estimated the fractions of tumor cells that exhibited these CNVs in each clone (**Fig. 1g**, **Supplementary Fig. 4a,** list of druggable genes from a personalized medicine trial ^20^. For instance, in the LFS_MB sample, targets such as *PD-L1, SMO* or *MYCN* were focally gained and highly expressed (using expression estimates from clone-matched scRNA-seq, see Methods and **Fig. 4** for details on the integration) in a subset or in all tumor cell clones in the primary tumor. However, in the matched relapse tumor LFS_MB1R, no druggable alteration would be prioritized for these three targets (**Fig. 1g**). While this highlights changes in the genetic landscape of the tumor, possibly as a response to treatment, our single-cell data enabled us to disentangle this variability at different target loci (**Fig. 1g, Supplementary Fig. 4a**). Considering inter-cell genetic heterogeneity and identifying cancer drivers and druggable genes present in each clone may have a major impact on treatment outcome in CT tumors. Importantly, both clonal (e.g. *PD-L1*) and subclonal targets (e.g. *SMO, MYCN*) were identified on CT chromosomes as well as within non-CT regions. Therefore, the heterogeneity in drug targets is only partially linked to chromothripsis heterogeneity and partially associated to CNVs independent of chromothripsis.

### Chromothripsis is a major event for the formation of double-minute chromosomes

Double-minute chromosomes, extrachromosomal circular DNA fragments carrying amplified oncogenes, were previously suggested to be generated by chromothripsis ^2,10^ but have not been studied at the single-cell level. In particular, the relationship between double-minute chromosomes and chromothripsis has not been systematically disentangled. The extent to which the double-minute chromosomes generated by chromothripsis themselves or the derivative CT chromosomes drive the selective advantage is unclear. To detect and quantify putative double-minute chromosomes, we searched for small fragments of a few megabases (Mb) highly amplified carrying oncogenes in the scDNA-seq data (**Fig. 2a**, sample MB243-Nuclei). By assembly from the matched bulk data, we could confirm that these amplified fragments were circular DNA structures (**Fig. 2b**). Interestingly, their copy number varied from 5 to more than 100 copies per nucleus in the single-cell data, suggesting an additional level of heterogeneity (**Fig. 2c**; Methods).

**Figure 2.**
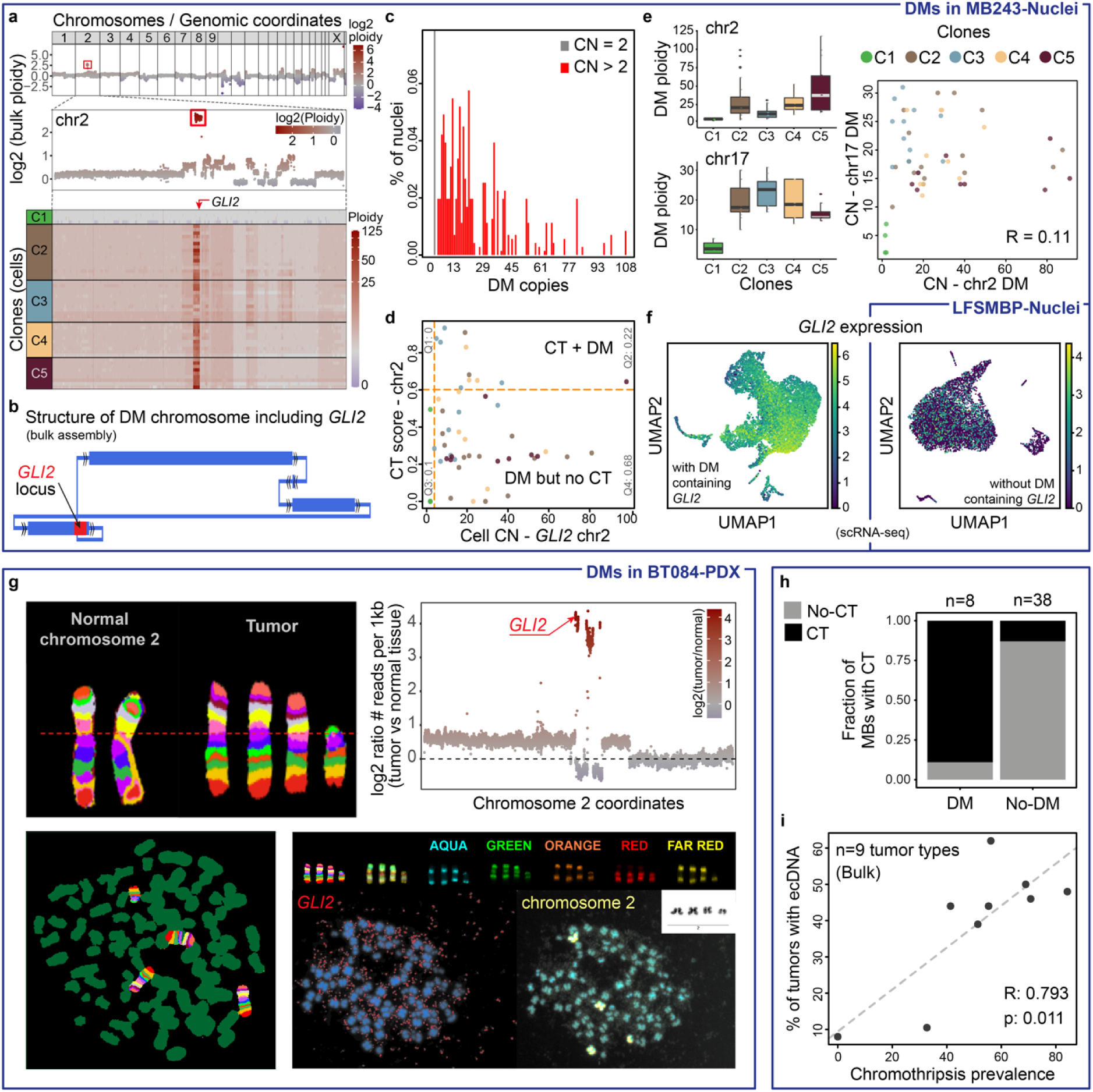
Chromothripsis is a major event for the formation of double-minute chromosomes. **a,** Representative example (MB243-Nuclei, shown in panels a to f) showing double-minute chromosomes carrying the *GLI2* oncogene (whole-genome view and zoom on chromosome 2) generated by a chromothriptic (CT) event. **b,** Structure of the double-minute chromosome carrying *GLI2* confirmed by bulk WGS assembly (Derived using AmpliconArchitect, a tool for reconstruction of ecDNA amplicon structures using WGS data; Methods). Blue rectangles show genomic segments. Arrows represent the orientation of a segment from lower to higher coordinates. **c,** scDNAseq of tumor nuclei shows high copy-numbers of the segments included in the double-minute chromosome. Most tumor cells carry 10 to 20 copies of the double-minute chromosome, with more than 100 copies per cell in extreme cases. **d,** Relationship between chromothripsis on chromosome 2 and the number of copies of double-minutes. Subsets of tumor cells carry both the CT chromosome and double-minute chromosomes while other cells keep the double-minute chromosome but have lost the CT chromosome. **e,** Number of copies of double-minute chromosomes per clone (left) and relationship between the number of copies of individual double-minute chromosomes present within one tumor (right). The data are represented as boxplots where the middle line corresponds to the median, the lower and upper edges of the box correspond to the first and third quartiles, the whiskers represent the interquartile range (IQR) ×1.5 and beyond the whiskers are outlier points. **f,** Tumor cells with double-minute chromosomes including *GLI2* show a high expression of this oncogene (left, sample MB243-Nuclei; right, *GLI2* expression in sample LFS_MBP-Nuclei for comparison with a sample without double-minute including *GLI2*). **g,** FISH validation of double-minute chromosomes carrying *GLI2* likely generated by chromothripsis on chromosome 2 (BT084-PDX sample). In addition to the *GLI2* probe (red), we used a multicolor probe for chromosome 2. **h,** Sonic Hedgehog medulloblastomas with double-minute chromosomes have a significantly higher chromothripsis prevalence. **i,** Chromothripsis is significantly linked with the presence of double-minute chromosomes across 9 tumor types.

By analyzing the relationship between CT chromosomes and double-minute chromosomes, we detected cells carrying both the CT chromosome and the double-minute chromosome (likely derived from the CT event), or only the double-minute chromosome but no CT chromosome (**Fig. 2d**). The presence of cells carrying only the double-minute chromosomes but without the derivative CT chromosome suggests that the double-minute chromosomes themselves may possibly provide a stronger selective advantage. The possible loss of the derivative CT chromosome in subsets of cells might explain the absence of direct correlation between the chromothripsis score and the number of double-minute chromosomes (**Fig. 2d**). In tumors with two or more double-minute chromosomes originating from distinct chromosomes, the amounts of double-minute chromosomes generated from different loci were tightly correlated in some tumors, but divergent in others, with nuclei characterized by the presence or absence of each individual double-minute chromosome (**Fig. 2e, Supplementary Fig. 4**). We detected substantial heterogeneity across clones but also across cells within clones in the number of double-minute chromosomes, suggesting that the number of double-minute chromosomes is not directly linked with the copy-number profiles (**Fig. 2e, Supplementary Fig. 4**). Essentially, this suggests that double-minute chromosomes further add to the intra-tumor genetic heterogeneity. Cells with extreme levels of double-minute chromosomes were rare and usually did not cluster with any other cell from the same tumor, suggesting that above a given threshold, the number of copies might not further increase the selective advantage for clonal expansion (**Supplementary Fig. 4**). The presence of double-minute chromosomes was associated with higher RNA expression of oncogenes carried on the double-minute chromosome, with large variations in expression within the tumor (**Fig. 2f**).

We experimentally validated the presence of double-minute chromosomes by FISH in the tumor BT084, focusing on the CT event on chromosome 2 (**Fig. 2g**). For this purpose, we combined a FISH probe for *GLI2*, an oncogene carried by double-minute chromosomes in this tumor, and a Xcyte 2 probe allowing us to visualize distinct regions of chromosome 2 with different fluorophores. The vast majority (close to 80%) of the analyzed metaphases showed four copies of chromosome 2, with three copies of similar size and one shorter copy. We occasionally detected metaphases with three or five copies of chromosome 2, consistent with the single-cell CNV data showing intra-tumor heterogeneity regarding the presence of the CT chromosome across cells mentioned earlier.

To further assess the link between chromothripsis and double-minute chromosomes in Sonic Hedgehog (SHH) medulloblastoma, we searched for double-minute chromosomes in bulk WGS of SHH medulloblastoma (n=48) and performed chromothripsis scoring (see Methods). Remarkably, medulloblastomas with double-minute chromosomes showed a significantly higher chromothripsis prevalence (**Fig. 2h**), confirming the association suggested in a previous study in one medulloblastoma via inference of the double-minute chromosome structure from bulk WGS ^2^. In addition, we showed a significant correlation between the presence of double-minute chromosomes and chromothripsis prevalence across nine tumor types (**Fig. 2i**). Hence, we hypothesize that chromothripsis is not only one way how double-minute chromosomes are generated, but might be the major way.

### Transcriptional heterogeneity in medulloblastomas with chromothripsis

Beyond selective advantages provided by chromothripsis, we set out to characterize further consequences of chromothripsis, in particular on the transcriptome of these tumors. Even though previous studies have considered scRNA-seq profiling to characterize medulloblastoma heterogeneity ^21,22^, the transcriptional consequences of a highly rearranged genome in CT medulloblastoma have not been described so far. We performed 10X single-cell (from PDX samples) and single-nuclei (from patient tumors) RNA-sequencing of the same 7 samples characterized on the genomic level (**Fig. 3a, Methods**). In total, we obtained 15,259 single-nuclei and 7,241 single-cell (PDX) transcriptomes after QC, respectively (hereafter referred to as scSeq for both cells and nuclei). Alignment across samples (**Supplementary Fig. 5a**) followed by analysis of literature-derived marker genes (**Methods**) identified 6 normal and 12 malignant cell types (**Fig. 3b-e**), which were also identified when analyzing individual tumors separately (**Supplementary Fig. 5,** see Methods). Considering the cells assigned to normal cell types, long-lived cells such as astrocytes and microglia were detected in the PDX samples as well as in the nuclei samples (**Fig. 3b-c, Supplementary Fig. 5b-c**). Other normal cell types such as endothelial cells, Purkinje cells or meninge cells were exclusively detected in nuclei sampled from primary tumors, suggesting that these normal cell types do not persist in a PDX model. The absence of specific cell types in PDX models has been previously reported in a different context (mentioned in ^23^ or in ^24^). Next, focusing on the subsets of malignant cells, we observed three major cell states that were present both in single-cell nuclei and scRNA-seq from PDX (**Fig. 3d-e**). The first malignant cell state, identified in all samples, was characterized by SHH signaling genes (e.g. *PTCH2, GLI2, HHIP*) as well as ribosomal genes and initiation/elongation factors. The second malignant cell state was characterized by proliferation and expression of markers of cell cycle activity and chromosome segregation (e.g. *MKI67, TOP2A, DIAPH3, HMGB, POLQ*). The third malignant state was characterized by neuronal development and differentiation markers (e.g. *RBFOX3, NEUROD1, CAMK2B, NCAM2*). Notably, the marker genes linked to all three cell states have been previously associated with non-CT SHH medulloblastomas ^21^. Consequently, despite pronounced genomic differences between CT and non-CT medulloblastomas from the same subgroup, our scRNA-seq data indicate no major difference at the level of transcriptional programs.

**Figure 3.**
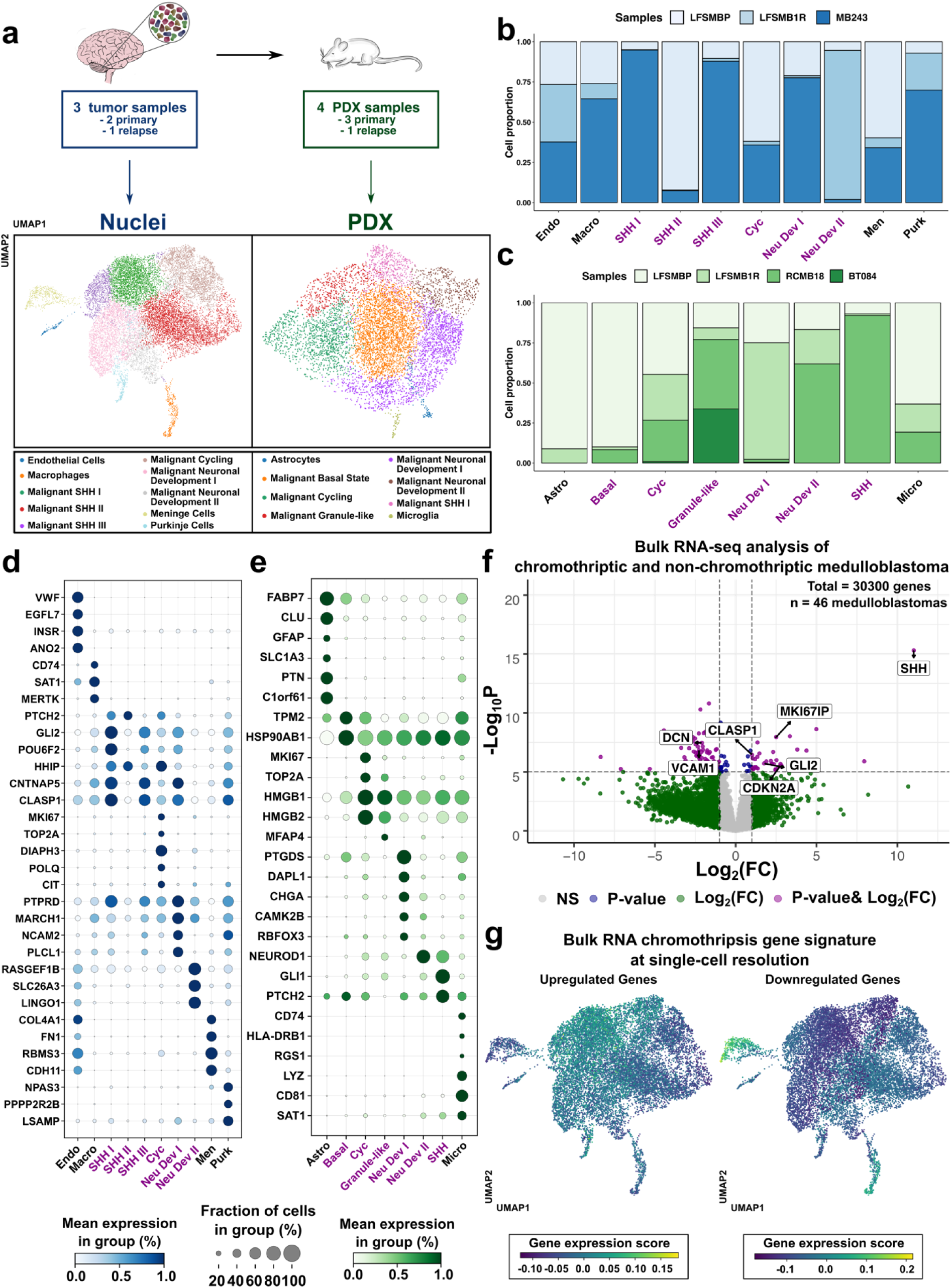
Distinct transcriptional programs dominate malignant cell types in medulloblastoma with chromothripsis. **a,** Overview of the experimental procedure to generate the single-cell and -nuclei RNA-seq data for the 7 samples of this study (Fig 1 and Fig 2). Left: single-nuclei RNA-sequencing (blue, tumors). Right: single-cell RNA-sequencing (green, PDX), based on two samples (LFS_MBP and LFS_MB1R) for which corresponding PDX models were available (**Supplementary Table 1**). UMAP embedding for tumor samples profiled using single-nuclei RNA-seq, (left, 15,259 cells) and PDX profiled using single-cell RNA-seq (right, 7,241 cells) from tumors and PDX (after quality control and alignment; **Methods**). Cell types annotated using literature-derived marker genes indicated in color (**Methods**). **b-c**, Stacked bar plots, displaying the relative prevalence of individual cell types across samples, either considering single-nuclei RNA-seq profiled tumors (**b**) or scRNA-seq profiled PDX models (**c**). Sample identifier encoded in color. **d-e,** Dotplot, displaying the expression level and prevalence of marker genes (x-axis) across the cell populations identified (y-axis) as in **a** and **b**. Dot size denotes the fraction of cells expressing the respective marker gene; dot color denotes the relative expression level. Malignant cell types highlighted in purple letters, non-malignant cell types shown in black. **f,** Volcano plot, displaying log2 fold change (x-axis) versus negative log p-values (y-axis) of genes identified from a differential expression analysis between chromothriptic (CT) and non-CT tumors (n = 46; multivariate analysis adjusting for tumor cell content and SHH group; **Methods**). Vertical and horizontal lines correspond to absolute log2 fold change values of 1 and p < 1E-5 respectively. **g,** UMAP embedding for tumor samples profiled with single-nuclei RNA-seq as in **a**, with cells colored by the expression signature of CT medulloblastomas derived from bulk differential expression analysis. Left: overexpression signature of CT medulloblastomas, right: underexpression signature of CT medulloblastomas.

In order to identify transcriptional signatures that are specific to CT tumors, we leveraged existing bulk RNA-seq data resources ^25^ (n=38 non-CT SHH medulloblastomas and n=8 LFS SHH medulloblastomas with CT, **Fig. 3f**, all tumors classified from SHH1 to SHH4). Differential expression analysis between CT and non-CT medulloblastoma identified 829 differentially expressed genes (FDR < 0.05, Log2FC < -1 & Log2FC > 1, Benjamini-Hochberg adjusted, two-sided Wald test, accounting for SHH subgroups, **Methods**). Genes associated with proliferation (e.g. *MKI67IP*), SHH signaling (e.g. *SHH*, *GLI2*), double-minute chromosomes (e.g. *CLASP1, GLI2)*, resistance (e.g. *FOXM1*) and MYC targets (e.g. *FKBP4, RRP1B*) were overexpressed in CT medulloblastomas compared to non-CT medulloblastomas (**Fig. 3f, Supplementary Fig. 6a**). In contrast, genes associated with differentiation (e.g. *DCN*, *VCAM1*) were expressed at lower levels in CT medulloblastoma compared to non-CT medulloblastoma. Gene Set Enrichment Analysis (GSEA, based on the top 2,000 up- and down-regulated genes) identified pathways including MTOR signaling and SHH signaling as significantly enriched in CT medulloblastomas (FDR < 0.05, **Supplementary Fig. 6b**).

To assess this expression signature in our single-nuclei dataset, we intersected the list of differentially expressed genes between LFS medulloblastomas with CT and non-CT SHH medulloblastomas with our scRNA-seq data and scored the relative activity of this signature (**Fig. 3g**; considering up- and down-regulated genes separately; scored relative to random reference gene sets; **Methods**). This identified heterogeneous activity of this signature in tumor-derived scNuclei, with malignant subpopulations being associated with genes upregulated in CT tumors, whereas differentiated cell populations were associated with downregulated genes (**Fig. 3g**). Notably, these differences were not detected in PDX samples (**Supplementary Fig. 6c**).

While the transcriptome analysis hinted towards minor differences between CT and non-CT tumors only, beyond transcriptional changes, DNA methylation signatures reflect features from the cell of origin, which may help to discriminate further between CT and non-CT tumors. Hence, to better understand the aggressiveness of CT medulloblastomas, we compared the DNA methylation patterns between both tumor subgroups (**Supplementary Fig. 6d**). After annotating the tumors by SHH subgroup (SHH1 to SHH4, n=1774) and *TP53* status, this suggested that CT medulloblastomas developing in LFS patients cluster together, between infant and children SHH medulloblastomas, and with SHH MBs harboring somatic *TP53* mutations. Altogether, the transcriptome and methylome profiles indicate more progenitor-like and undifferentiated cells as compared to non-CT tumors, which display a less pronounced stem-like signature.

### Integration of copy number clones from scDNA with scRNA-seq data reveals the connection between genome instability and transcriptome

Having detected distinct malignant and non-malignant cell states, we set out to evaluate how the somatic diversity between clones is linked to transcriptional differences. Established methods such as clonealign ^26^ have been proposed to identify clonal populations in independent scDNA- and scRNA-seq data. However, this method is not designed for 10X Genomics data and hence we developed an alternative approach which relies on the estimation of relative CNV profiles based on scSeq data using inferCNV ^27^ (using a detected normal cell population as reference, **Methods**). Applying inferCNV to our data yielded CNV profiles that were qualitatively consistent with CNV profiles obtained from scDNA-seq (**Supplementary Fig. 7**). Next, we leveraged these parallel CNV profiles in both scDNA- and scRNA-seq modalities to align single transcriptomes to the scDNA-seq clones described earlier (**Fig. 1**). Briefly, individual transcriptomes were assigned to the clone with the closest matching CNV profile (**Fig. 4a**, **Supp. Fig. 8a**; using permutations to derive an empirical measure of alignment confidence; **Methods**).

**Figure 4.**
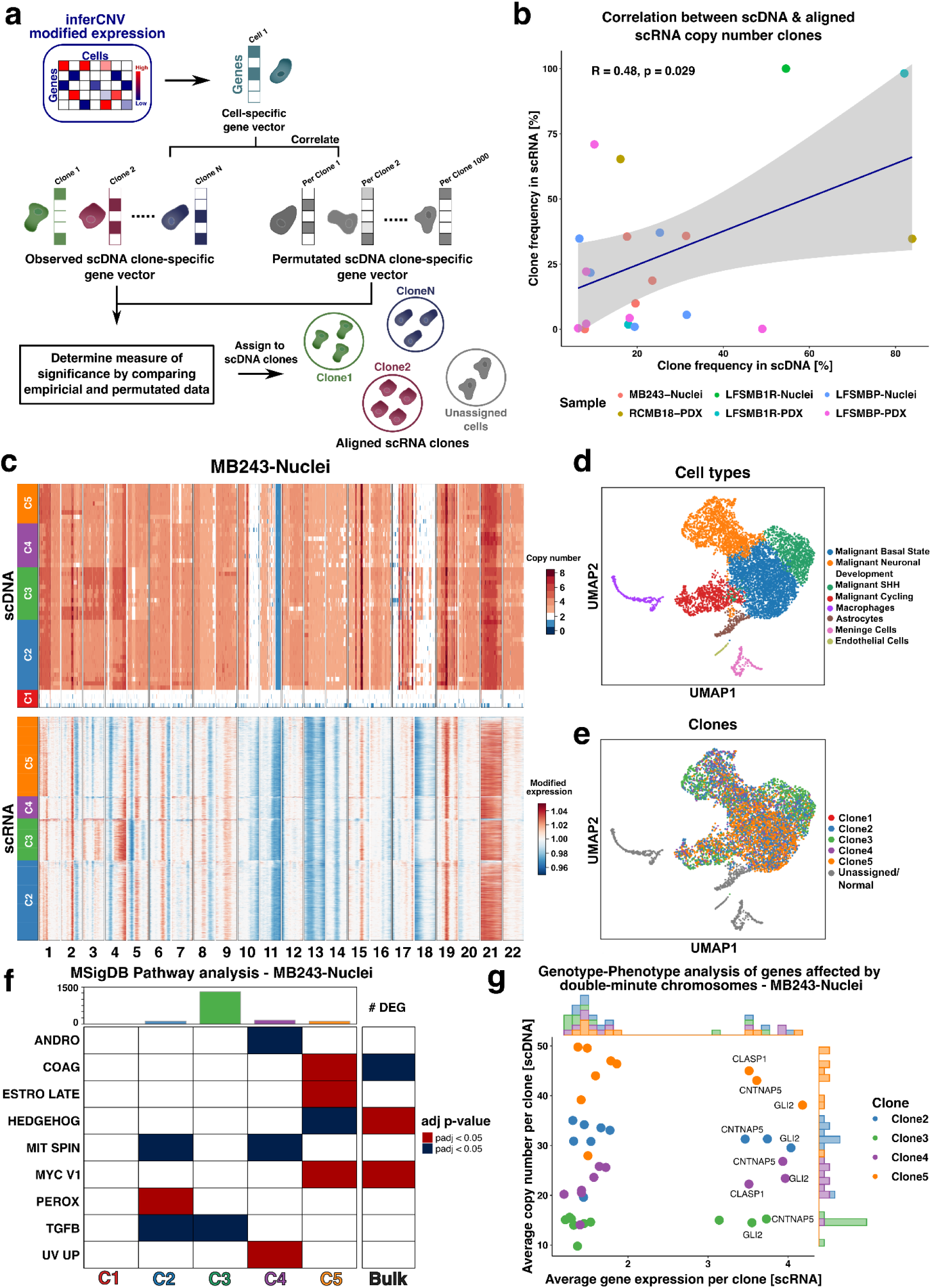
Integrating DNA and RNA on single cell level reveals copy number related pathway alterations. **a,** Schematic of the computational strategy for aligning scRNA-seq profiles to CNV profiles of clones derived from scDNA-seq. **b,** Scatter plot between the clonal fraction estimated from scDNA-seq versus the fraction of aligned scRNA-seq profiles. The blue line denotes line regression fit with shaded areas indicating the 95% confidence interval (grey area). Color denotes the sample of origin. **c,** Results from the clone alignment of scDNA- and scRNA-seq profiles for the sample MB243. Top: Heatmap displaying the copy number profiles derived from scDNA, with color corresponding to copy number (scale [0,8]; larger values clipped). Bottom: Heatmap showing relative CNV profiles estimated using inferCNV ^27^, with color corresponding to modified expression (scale centered on 1; diploid state; Methods). **d,** UMAP embedding depicting 7,302 cells from scRNA-seq after QC for MB243-Nuclei. Cells are colored according to their assigned cell type identity. **e**, Same UMAP as in **d**, with cells colored according to the clones from the integrated analysis described in **a**. Phenotypically normal cells (n = 606 cells) as well as cells that did not meet the assignment confidence (Bonferroni adjusted p > 0.05 **Methods**) were excluded and are marked as unassigned / normal cell population (grey). **f,** Heatmap, displaying enriched pathways across clones for MB243-Nuclei and bulk RNA-seq. The number of clone-specific differentially expressed genes (FDR < 0.05, two-sided Wilcoxon rank sum test, Benjamini-Hochberg adjusted, one clone versus all) is shown as a bar plot on the top. Each bar is colored based on the clone color shown in **c** and **e**. For the pathways, significance is shown as p < 0.05 (Kolmogorov–Smirnov statistic, Benjamini-Hochberg adjusted). Depending on the type of differential expression (up- or down-regulation), the pathways are shown in shades of red or blue. Identical pathways identified from differential expression analysis between CT and non-CT tumors from bulk RNA-seq shown for reference. **g,** Scatterplot representing the genotype to phenotype analysis of genes included in double-minute (DM) chromosomes for MB243-Nuclei (see Methods). The mean copy number value per clone in scDNA (y-axis) and the mean normalized gene expression per aligned clone from scRNA (x-axis) are shown. Each dot represents a gene colored according to its clone identity. The histograms show the distribution of expression and copy number values, evenly colored according to the clones. ANDRO: androgen receptor signaling, COAG: Coagulation, ESTRO LATE: Estrogen response late, HEDGEHOG: Sonic Hedgehog signaling, MIT SPIN: mitotic spindle, MYC V1: MYC pathway signaling, OXPHOS: oxidative phosphorylation, PEROX: peroxisome, TGFB: TGFbeta signaling, UV UP: UV signaling upregulated.

Across all samples, this strategy allowed for assigning a clone identity to between 46% and 100% of single-cell transcriptomes (**Supplementary Table 3**), which resulted in broadly consistent clone frequency estimates between the scDNA data and the integrated clone frequency in scRNA-seq (**Fig. 4b**, R^2^ = 0.48, p = 0.029, linear model). Focusing on MB243 with the largest number of aligned cells (**Fig. 4c**; **Supplementary Fig. 8b**; 100% alignment; n = 7,302 scRNA-seq transcriptomes), the observed mixing of clone labels and cell types indicates that the dominant source of variation in the scRNA-seq data is cell type rather than clone (**Fig. 4d-e, Supplementary Fig. 9**). Next, we used the clone assignment to identify differentially expressed genes and pathways between clones (**Fig. 4f**, FDR < 0.05, two-sided Wilcoxon rank sum test, Benjamini-Hochberg adjusted, one clone versus all). Pathway analysis of DE genes identified, among others, significant up-regulation of MYC targets and down-regulation of Hedgehog signaling in clone C5, in line with the differential expression analysis between CT and non-CT medulloblastomas (**Fig. 3f, Supplementary Fig. 6**) and TGFbeta signaling in clones C2 and C3 (FDR < 0.05, Kolmogorov–Smirnov statistic, Benjamini-Hochberg adjusted). Repeating the analogous analysis across all samples (**Supplementary Fig. 8**) identified recurrent clone-specific up-regulation of genes related to G2M checkpoint, the mitotic spindle or E2F targets, which notably mirrors the pathways identified between CT and non-CT medulloblastoma (**Fig. 4f**, **Supplementary Fig. 8).** Given the link with *TP53* deficiency and the importance of dysfunction of the cell division for chromothripsis, this added further confidence in our integration approach. Furthermore, the clone alignment revealed a bimodal expression pattern of genes located on the double-minute chromosomes in MB243, with only a subset of genes showing a strong overexpression (**Fig. 4g**). Interestingly, *GLI2* and *CLASP1,* essential players in SHH signaling and in microtubule dynamics, were highly overexpressed in the bulk RNA-sequencing analysis (**Fig. 3f**), suggesting a major role in CT tumors. Taken together, the results from our integration suggest that the combined information from single-cell genomes and transcriptomes can reliably identify copy-number related pathway alterations, which are biologically relevant. Future studies focusing on larger cohorts of CT tumors will be crucial to further underline these findings and highlight the impact of copy number variation on the transcriptome.

### Loss of chromosome 3p and *SETD2* deficiency as early events potentially facilitating chromothripsis

In addition to characterizing pathways activated in specific clones, we leveraged the single-cell and bulk WGS data to identify putative early alterations that might contribute to chromothripsis occurrence. Previous studies indicated that inactivation of essential checkpoints likely facilitates chromothripsis and/or the survival of a cell after CT events ^2,11,12^. We searched for early events potentially linked with chromothripsis. Loss of chromosome 17p (chr17p), carrying the wild-type *TP53* allele, was already known to be associated with chromothripsis in medulloblastoma patients with germline *TP53* mutations ^2^. In agreement with this, rare non-tumor cells with a balanced profile (defined as non-tumor cells based on the absence of CNV), except a focal loss of the *TP53* locus, supported the loss of p53 as an early event in CT tumors (**Supplementary Fig. 10a**). In addition, we identified chromosome 3p (chr3p) loss as a clonal event linked with chr17p loss and chromothripsis in our single-cell data (**Fig. 5a-b**). This was further supported by phylogenies reconstructed from deep bulk sequencing data and allele frequency analyses (**Fig. 5c**). Investigating this association in a cohort of 227 medulloblastomas, we found that loss of chr3p was highly significant when searching for genomic regions tightly linked with chromothripsis (**Fig. 5d-e**, two-sided Fisher exact test, p<10^-5^ and two-sided Chi-square test, p<10^-8^, respectively). Importantly, loss of chr3p was also significantly linked with chromothripsis in breast and lung cancer (two-sided Chi-square test, p<1.32×10^-4^ for breast cancer and p<3.44×10^-10^ for lung cancer), suggesting a potential pan-cancer relevance, beyond medulloblastoma (**Supplementary Fig. 10b**). To validate this association experimentally, we performed time-course analyses with primary cells from LFS patients. In these primary cultures, we identified loss of both chromosomes 17p and 3p as early events linked with chromothripsis using WGS (**Supplementary Fig. 10c**).

**Figure 5.**
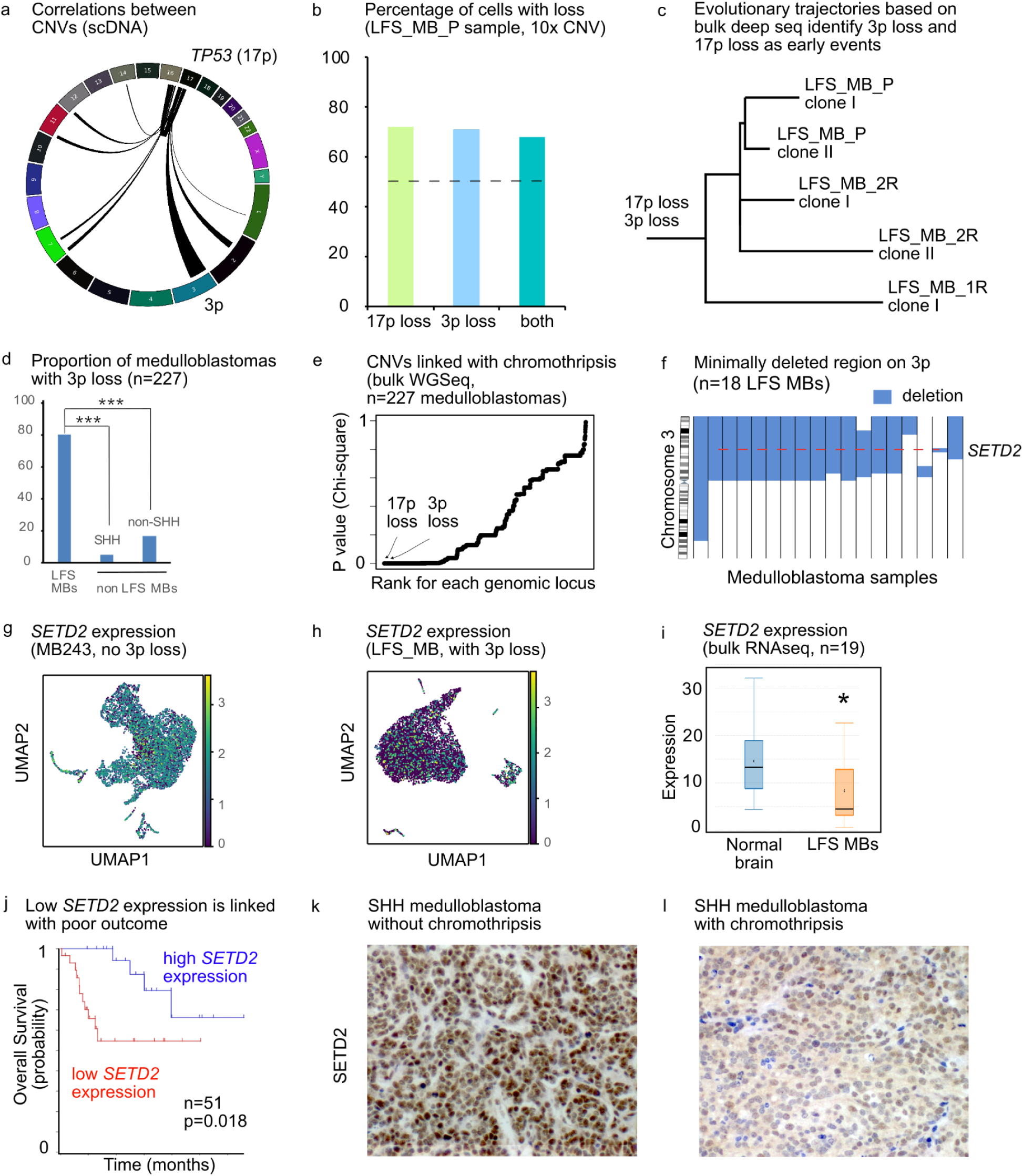
Combining single-nuclei DNA-seq and bulk whole-genome sequencing identifies early events potentially facilitating chromothripsis. **a,** CNVs correlation analysis from single-cell DNA sequencing data identifies 3p loss as an event linked with *TP53* loss on 17p. Arcs between chromosomes indicate regions with correlated CNV profiles (two-sided correlation test, Pearson r>=0.77, p < 0.01, Bonferroni adjusted). **b,** Proportion of nuclei with 17p loss, 3p loss or both in the LFS_MBP sample. The proportion of nuclei with loss of both 17p and 3p is above the dashed line indicating what would be expected by chance. **c,** Evolutionary trajectories based on deep WGS identify 17p loss and 3p loss as early events (longitudinal analysis of three matched tumor samples, ST). **d,** Proportion of medulloblastomas with 3p loss among tumors with chromothripsis (LFS medulloblastomas) and non-LFS medulloblastomas. Two-sided Fisher exact tests were performed to compare the proportions of tumors with 3p loss between LFS medulloblastomas and non-LFS medulloblastomas. **e,** 17p loss and 3p loss are significantly linked with chromothripsis in medulloblastoma (bulk WGS, n=227 medulloblastomas). f, *SETD2* is located in the minimally deleted region on 3p in medulloblastoma. Blue shades show the size and location of the deleted region. Each vertical bar shows one medulloblastoma with chromothripsis. **g, h,** Loss of 3p leads to decreased *SETD2* expression (scRNAseq). **i,** Loss of 3p leads to decreased *SETD2* expression (bulk RNAseq; the central line shows the median, minimum and maximum values are shown by whiskers). Statistical significance was tested using one-tailed t-test. **j,** Low *SETD2* expression is linked with poor survival in medulloblastoma (log-rank test, SHH alpha subgroup). **k, l,** Representative examples of medulloblastomas with or without chromothripsis showing low or high SETD2 protein expression, respectively.

To search for candidate genes on chr3p potentially preventing chromothripsis, we defined the minimally deleted region across bulk data from 20 LFS medulloblastomas (**Fig. 5f**). We narrowed the list of candidates based on gene expression in LFS medulloblastomas, reported mutations and function. Among the evaluated genes, *SETD2* was a promising candidate, due to the known tumor suppressive role of the SETD2 methyltransferase, lost or mutated in various cancers, and its importance for DNA replication, DNA repair and genome instability ^28,29^. Medulloblastomas with chr3p loss displayed a lower *SETD2* expression based on single-cell and bulk RNAseq (**Fig. 5g-i**). In addition, low *SETD2* expression was linked with significantly shorter overall survival in SHH medulloblastoma (**Fig. 5j**, two-sided log-rank test, p<0.02, SHH alpha (SHH3) subgroup, which is the molecular subgroup to which most CT medulloblastomas belong). Sonic Hedgehog medulloblastomas with chromothripsis displayed a lower protein expression as compared to SHH medulloblastomas without chromothripsis, as shown by immunohistochemistry (**Fig. 5 k-l**).

As combined single-cell and bulk sequencing analyses identified *SETD2* as a promising candidate potentially preventing chromothripsis, we analyzed the functional consequences of *SETD2* loss. To test for a potentially causal role of *SETD2* in chromothripsis, we used CRISPR/Cas9 to inactivate *SETD2* in p53 wild-type and p53-deficient neural stem cells, respectively (**Fig. 6a**). Chromothripsis has previously been linked with genome doubling ^30^, as well as with the formation of micronuclei ^13,17^, which are abnormal nuclear structures containing one or very few chromosomes. Our CRISPR/Cas9 experiments showed that, upon *SETD2* inactivation in a p53-deficient background, the formation of micronuclei significantly increased as compared to inactivation of *TP53* only (**Fig. 6 b-c**, one-way Anova and Bonferroni multiple comparison tests, p<0.05). In addition, as compared to wild-type cells, *TP53*/*SETD2* knock-out cells showed a significantly larger nuclear area (**Fig. 6d**, one-way Anova and Bonferroni multiple comparison tests, p<0.05), a measure which is used as a surrogate marker for polyploidization ^31^. Immunofluorescence analysis of the commonly used DNA double-strand break marker γH2AX showed a significant increase in the levels of DNA double-strand breaks in *TP53*/*SETD2* knock-out cells as compared to wild-type neural stem cells (**Fig. 6 e-f**, one-way Anova and Bonferroni multiple comparison tests, p<0.05). Double stain for phosphorylated histone H3 and acetylated tubulin identified aberrant mitoses in *SETD2* and in *TP53*/*SETD2* knock-out cells, such as failure to congress at prometaphase, multipolar spindle formation and anaphase bridges (**Fig. 6g**). This is in agreement with chromothripsis being one consequence of bridge breakage ^16^. Finally, we measured a significantly increased proliferation rate upon *SETD2* inactivation of *TP53* and *SETD2* (**Fig. 6h**), indicating a selective advantage. Altogether, the functional consequences of the inactivation of *TP53* and *SETD2* in neural stem cells suggest a possible causative role for these two genes in the occurrence of chromothripsis in medulloblastoma.

**Figure 6.**
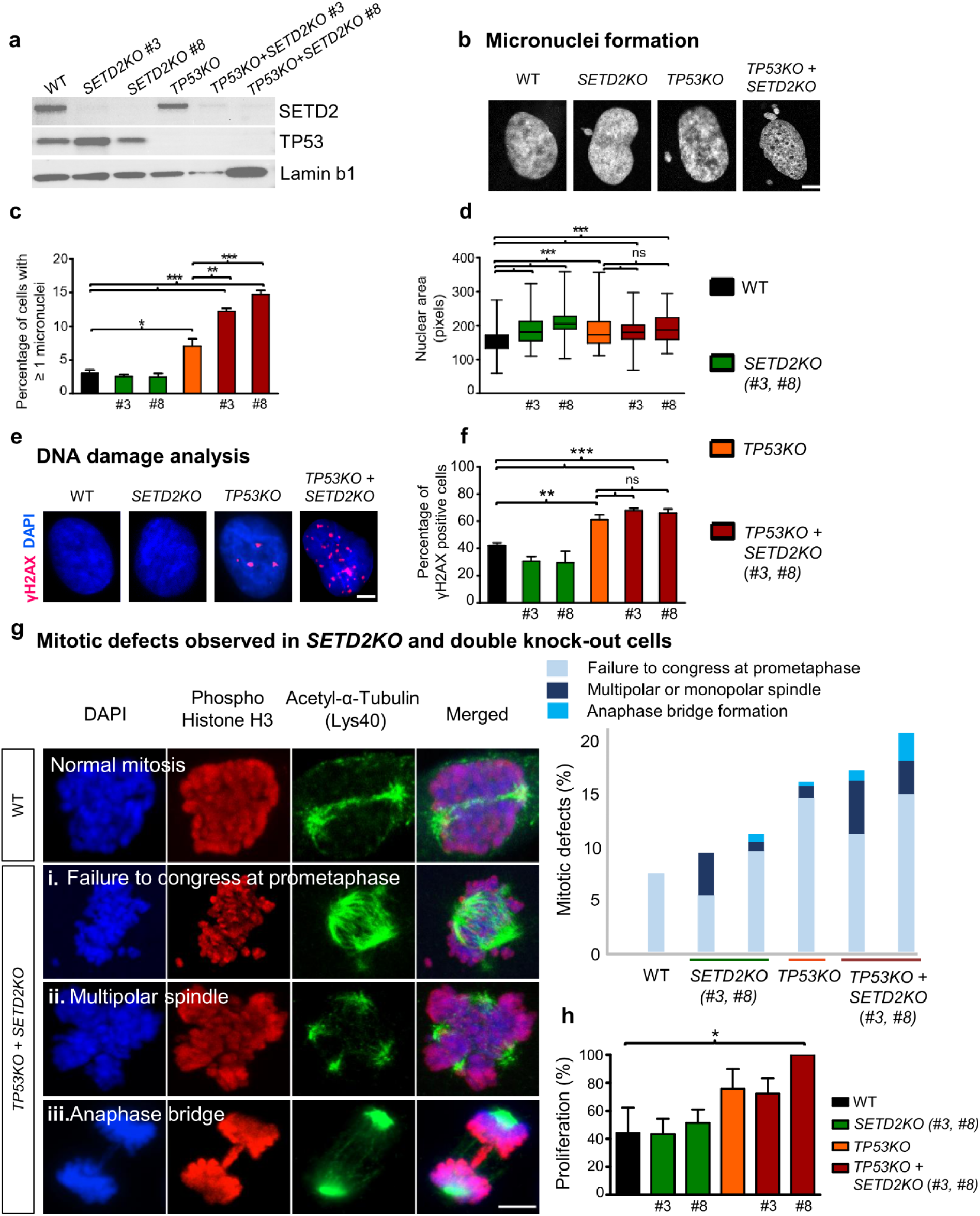
Inactivation of *SETD2* and *TP53* in neural stem cells leads to genome instability. **a,** Western blot analysis confirms efficient inactivation of *SETD2* and *TP53* in neural stem cells by CRISPR/Cas9. Two independent guide RNAs were used for *SETD2* (guides #3 and #8). **b,** Inactivation of *SETD2* in a p53 deficient background leads to the formation of micronuclei, aberrant nuclear structures linked with genome instability. Representative pictures based on three independent experiments are shown. Scale bar, 5 µm **c,** Quantification of micronuclei (three independent experiments; mean±SEM). **d,** Inactivation of *SETD2* leads to a larger nuclear area (three independent experiments; the central line shows the median, the minimum and maximum values are displayed with whiskers). **e, f,** Inactivation of *SETD2* in a p53 deficient background leads to high levels of DNA double-strand breaks. Immunofluorescence analysis of γH2AX foci and quantification of γH2AX positive cells (five independent experiments; mean±SEM). **g,** Inactivation of *SETD2* in a p53 deficient background leads to mitotic defects, as shown by immunofluorescence analysis of phospho histone H3 and acetyl tubulin. **h,** Inactivation of *SETD2* in a p53 deficient background leads to increased proliferation rate. Metabolic activity is shown as a percentage of the sample with the highest values (five independent experiments; mean±SEM). One-way ANOVA and Bonferroni multiple comparison tests were used to test for statistical significance in **c**, **d**, **f** and **h**.

## DISCUSSION

Li-Fraumeni syndrome (LFS) medulloblastoma is a clinically challenging type of childhood brain tumor, where patients suffer from a dismal prognosis. These tumors are characterized by chromothripsis, an extreme phenomenon of genome instability, which is present in close to 100% of these medulloblastomas ^2,19^. Hence, understanding the genomic heterogeneity and its consequences on the transcriptome are essential to identify targets for novel therapeutic strategies for this subgroup of patients.

In this study, we combined single-cell analyses of the genome and transcriptome together with bulk deep sequencing to provide a roadmap of alterations in CT medulloblastoma with *TP53* germline mutations. By combining bulk and single-cell DNA sequencing of matched tissue samples, we scored chromothripsis events at clonal resolution. This approach enabled us to shed light into the genomic heterogeneity at the level of CT chromosomes, cancer drivers as well as potentially druggable targets. In addition, we observed and experimentally validated an association between the abundance of double-minute chromosomes and chromothripsis, further increasing the bespoken heterogeneity. Comparisons between matched primary and relapse samples in patient tumors and PDX, on both genome and transcriptome, supported substantial heterogeneity with major implications for treatment. Importantly, our results also question the common view of chromothripsis as a single early event in tumor development, which goes along with limited intra-tumor heterogeneity.

We aimed at identifying putative early events in LFS tumor evolution. It has been unclear whether the inactivation of essential checkpoints such as p53 and others, may occur shortly before or after chromothripsis. Here, our results highlighted chr3p loss and *SETD2* inactivation as a potential early event facilitating chromothripsis occurrence. We experimentally underlined this observation utilizing CRISPR/Cas9-mediated inactivation of *SETD2* in p53 wild-type and p53-deficient neural stem cells. In line with this, we detected rare non-tumor cells with *TP53* loss, but no tumor clones with loss of chr17p and/or chr3p without chromothripsis. However, as such tumor cells are expected at a very low frequency, sequencing thousands of tumor cells would be necessary to detect such rare populations. To recapitulate the sequence of events, we used a time-course experiment, culturing primary fibroblasts from early passages with stable copy-number profiles to late passages with spontaneous chromothripsis occurrence. Our findings validated chr17p loss and chr3p loss as early events linked with chromothripsis.

So far, the transcriptional consequences of chromothripsis in tumors have not been systematically investigated. Here, we leveraged single-cell and single-nuclei RNA-seq to analyze 7 samples from LFS medulloblastoma and PDX samples. Remarkably, we found a variety of malignant and non-malignant cell types, a subset of which were represented in the PDX samples. Furthermore, we observed three transcriptional programs largely defined by i) SHH genes, ii) proliferation and iii) genes implicated in neuronal development. These programs were surprisingly consistent with programs previously observed in non-CT SHH medulloblastoma ^21^. Our analysis of bulk RNA-seq from CT and non-CT medulloblastomas emphasized differences in gene expression and activated pathways, in particular regarding stemness, resistance, SHH signaling and proliferation. However, future studies focusing on the origin of the aggressiveness of CT tumors will be needed in order to dissect the precise mechanisms explaining the poor outcome of patients with CT tumors.

Lastly, to link the genome and the transcriptome data, we demonstrated how copy number estimates allow for assigning single-cell transcriptomes to individual CNV clones. Reassuringly, our method identified the copy number clones from scDNA at a similar proportion in the scRNA-seq data, represented by a significant correlation between the frequency of clones in both data modalities. By linking distinct transcriptional profiles to the identified tumor clones, we were further able to highlight activated pathways, including but not limited to SHH signaling and MYC targets, in line with the DE pathways between CT and non-CT tumors, but also G2M checkpoint genes, the mitotic spindle apparatus and E2F targets. In the context of germline *TP53* deficiency, these results are not surprising, as loss of p53 allows cells to proceed through the cell cycle with mis-segregated chromosomes, eventually contributing to chromothripsis.

This study does not come without limitations. A larger sample size would presumably be needed in order to identify commonalities between chromothripsis and the establishment of double-minute chromosomes. In addition, although we investigated potential druggable targets in both scDNA and scRNA data, a larger number of matched primary tumors and relapse samples would be required to understand the influence of treatment on the intra-tumor heterogeneity in LFS medulloblastoma. Following this further, we would envision a larger cohort of CT and non-CT medulloblastomas being essential to get insights into the origin of the poor prognosis and aggressiveness of these tumors.

Tumors in LFS patients constitute a paradigm for the understanding of chromothripsis. Our work focusing on this group of patients can provide a roadmap from where the findings may be extended to different contexts, as the link between chromothripsis and *TP53* mutations also holds true outside the context of constitutive defects (e.g. in prostate cancer ^32^ or breast cancer ^33^. In the future, a more refined single-cell landscape of CT tumors will be needed to further confirm and increase the understanding of the genomic heterogeneity, diversity of cell types and active transcriptional programs. Unraveling the extent of genomic heterogeneity will be necessary to detect actionable targets, determine the evolutionary history and defeat the evolutionary capacity of tumor cells with high genome instability.

## METHODS

### Experimental Methods

#### Sample cohort, DNA extraction and whole genome sequencing

Human clinical samples and data were collected after receiving written informed consent in accordance with the Declaration of Helsinki and approval from the respective institutional review boards. All tumors used for bulk sequencing had a tumor cell content confirmed by neuropathological evaluation of the hematoxylin and eosin stainings. DNA was extracted from frozen tissue using Qiagen kits. Purified DNA was quantified using the Qubit Broad Range double-stranded DNA assay (Life Technologies, Carlsbad, CA, USA). Genomic DNA was sheared using an S2 Ultrasonicator (Covaris, Woburn, MA, USA). Whole-genome sequencing and library preparations for tumors and matched germline controls were performed according to the manufacturer’s instructions (Illumina, San Diego, CA, USA or NEBNext, NEB). The quality of the libraries was assessed using a Bioanalyzer (Agilent, Stockport, UK). Sequencing was performed using the Illumina X Ten platform.

#### Sample collection and establishment of patient derived xenografts

Orthotopic patient-derived xenografts were done in 6-10-week-old female immune-compromised mice (NSG, NOD.*Cg-Prkdc^scid^Il2rg^tm1Wjl^*). Patient-derived tumor cells were injected into the cerebellum, as described previously ^34^. All animal experiments were performed in accordance with ethical and legal regulations for animal welfare and approved by the governmental council (Regierungspräsidium Karlsruhe, Germany).

#### Nuclei isolation from tumor tissue

Frozen tumor tissue was used for nuclei isolation. Tissue was cut using a scalpel with 1mL of lysis buffer. After adding 4mL of lysis buffer, the suspension was transferred to a glass douncer. A total of 20 strokes were used to dounce the suspension on ice with two different types of pestles. The entire suspension was then filtered with a 100 µM filter and then with a 40 µM filter into precooled falcon tubes. After centrifugation for 5 min at 555 g at 4°C, the supernatant was removed and the pellet was resuspended in 5 mL of lysis buffer without Triton-X and DTT. This centrifugation step and the resuspension were carried out 3 times in total. The final pellet was then resuspended in 1mL of nuclei storage buffer in 1.5 mL LoBind Eppendorf tubes for further analysis.

#### Cell suspension from Patient-Derived Xenograft (PDX) cells

Viably frozen cells were thawed in a 37°C water bath followed by suspension in high purity grade PBS supplemented with 10% fetal calf serum. Cells were washed twice with PBS and stained with propidium iodide to select for viable cells. Cells were then filtered (40 µm) and sorted.

#### FACS

Cell suspensions from PDX models were sorted on a FACSAria to exclude dead cells and cell doublets. Contaminating mouse cells were excluded based on the side forward scatter.

#### 10X single-cell RNA-sequencing library preparation

The single cell suspensions of PDX cells or nuclei from frozen tissue specimens were loaded on a 10x Chromium Single Cell instrument (10x Genomics, California) to generate single-cell Gel Bead-In-Emulsions (GEMs). Single-cell RNA-Seq libraries were prepared using Chromium Single-Cell 5′ Library and Gel Bead Kit (PN1000014, 10x Genomics). Barcoding and cDNA synthesis were performed according to the manufacturer’s instructions. In short, GEMs were created where all cDNA from one cell shared a common 10x barcode. GEMs were then incubated at RT and cleaned up using Dynabeads. After post GEM-RT clean-up, full length cDNA was generated by PCR with a total of 14 cycles for library construction. The cDNA libraries were constructed using the 10x ChromiumTM Single Cell 5’ Library Kit according to the manufacturer’s protocol. In brief, the major steps for the library preparation included i) Target enrichment from cDNA, ii) Enriched library construction, iii) 5’ Gene expression library construction and QC. For final library QC, 1 ul of the sample was diluted 1:10 and ran on the Agilent Bioanalyzer High Sensitivity chip.

#### 10X single-cell DNA-sequencing library preparation

The single-cell suspensions from tumor nuclei or PDX cell samples were processed using the Chromium Single-Cell CNV Kit (10× Genomics) according to the manufacturer’s protocol. In brief, using cell bead polymer, single cells or nuclei were partitioned in a hydrogel matrix on Chromium Chip C. Once the cell beads were encapsulated and incubated, they were subjected to enzymatic and chemical treatment. This lysed the encapsulated cells and denatured the gDNA in the cell bead, to make it accessible for further amplification and barcoding. A second encapsulation was performed to achieve single cell resolution by co-encapsulating a single cell bead and a single barcoded gel bead to generate GEMs. Immediately after GEM generation the gel bead and cell bead were dissolved. Oligonucleotides containing standard Illumina adaptors and 10x barcoded fragments were then amplified with 14 PCR cycles during two-step isothermal incubation. After incubation the GEMs were broken and pooled 10x barcoded fragments were recovered. For final sequencing library QC, 1ul of the sample was diluted 1:10 and ran on the Agilent Bioanalyzer High Sensitivity chip. Although the experiment was performed, the library for the BT084-PDX sample did not pass the quality control steps and hence was not included in this study.

#### Sequencing of single-cell DNA and RNA libraries

Single-cell libraries were sequenced on the Illumina NextSeq and NovaSeq (paired-end sequencing).

#### Fluorescence in situ hybridization (FISH)

Nick translation was carried out for BAC clones all obtained from Source Bioscience (*GLI2*, clone RP11 297J22). FISH was performed on metaphase spreads using fluorescein isothiocyanate-labeled probes and rhodamine-labeled probes. Pre-treatment of slides, hybridization, post-hybridization processing and signal detection were performed as described previously. Samples showing sufficient FISH efficiency (>90% nuclei with signals) were evaluated. Signals were scored in, at least, 100 non-overlapping metaphases. Metaphase FISH for verifying clone-mapping position was performed using peripheral blood cell cultures of healthy donors as outlined previously. After the *GLI2* FISH to detect double-minute chromosomes, the coverslip was removed, and metaphase spreads were washed. Denaturation was performed to remove the signal from the *GLI2* probe and hybridization was done using the multicolor XCyte 2 probe from Metasystems according to the manufacturer’s instructions.

#### Cell culture

Neural stem cells (human iPSC derived NSCs, kindly provided by Dr. Daniel Haag) were cultured in matrigel (Corning, 356230) coated 6-well plates, in NeuroCult NS-A proliferation media kit (Stemcell Technologies, 05751) supplemented with 40 ng/ml EGF (Sigma, E4127), 40 ng/ml FGF (Preprotech, GMP100-18 B), 10 ng/ml hLIF (Millipore, LIF1010) and 10 µM Rock inhibitor Y-27632 (Enzo, ALX-270-333-M001).

#### CRISPR-Cas targeted gene disruption

Guide RNAs for *TP53* and *SETD2* were constructed and cloned into lenti CRISPR v-2 (Addgene, 52961) according to the original online protocol of the Zhang lab (http://www.genome-engineering.org/crispr/wp-content/uploads/2014/05/CRISPR-Reagent-Description-Rev20140509.pdf). Following genes were targeted:

*TP53* (gRNA2:CGACCAGCAGCTCCTACACCGG)

*SETD2* (gRNA3:AATGAACTGGGATTCCGACG)

and (gRNA8:GGACTGTGAACGGACAACTG).

Virus production and transduction were done as described previously; in brief, pLentiV2, pDMDG.2, and pSPAX were co-transfected in HEK293T cells, and virus-containing supernatant was concentrated by ultracentrifugation. Transduction was done by adding concentrated virus particles to NSC lines for 24 h, after which cells were maintained under selection either with puromycin or blasticidin at 2 µg/mL for 2-4 weeks. For CRISPR-mediated disruption of *TP53*, an additional selection for functional knockout was done using 20 µM nutlin treatment. After selection, cell lysates were made for western blotting or cells were grown for further experiments.

#### Western blotting

For western blot experiments, NSCs were detached with accutase (Sigma, A6964), collected in media and washed three times with ice-cold PBS. The pellet was resuspended and incubated for 10 min on ice in RIPA buffer containing Complete™, EDTA-free Protease Inhibitor Cocktail (Sigma, 4693159001) and benzonase (Millipore, 71205-3). Protein concentration was estimated using BCA assay. Lysates were mixed with 10% DTT bromophenol blue and boiled at 95 °C for 5 min. A total amount of 30 µg protein was loaded per lane of NuPAGE Tris-Acetate Protein Gel 3-8% (Life Technologies, EA0375BOX) and separated according to the manufacturer’s instructions. Immunoblotting was done on PVDF membranes in a tank blot system, using a borate-based buffer system (25 mM sodium borate, 1 mM EDTA, pH 8.8). Membranes were blocked with 5% milk powder in TBST for 1 h and probed with SETD2 (Cell signaling, 80290) 1:1000 overnight at 4°C with agitation, TP53 (Santa Cruz, sc-126) 1:500 and lamin b1 (Abcam, ab16048) 1:500 for 1 hour at room temperature. Membranes were washed with TBST and incubated for 30 min with HRP coupled secondary anti-mouse or anti-rabbit antibodies (Dianova, 115-035- 003 and 211-032-171) 1:5000. After washing, detection was done using enhanced chemiluminescence and images were recorded with Bio-rad Imaging System (LI-COR Biotechnology).

#### H&E stain and immunofluorescence

Hematoxylin and eosin (H&E) stain and immunofluorescence stain were performed on 4 µm formalin-fixed paraffin-embedded sections. Sections were deparaffinized, antigen retrieval was performed in 10 mM citrate buffer pH 6.0 for 40 min and sections were cooled down to room temperature. H&E stain was evaluated by a neuropathologist. For immunohistochemistry, rabbit SETD2 antibody (Atlas, HPA042451) was used at 1:500 with the DCS SuperVision 2 HRP Kit.

#### Immunofluorescence analysis of γH2AX in neural stem cells

Cells grown on coverslips were washed with PBS and incubated 15 min in 4% formaldehyde (formalin solution buffered at pH 6.8, Merck). Cells were washed once with 50 mM ammonium chloride (Carl Roth) and twice with PBS before permeabilization with 0.1% triton (Triton X-100, Gerbu Biotechnik). Blocking was done with 10% donkey serum (Merck). Primary antibody (S139, Abcam, 1:200 in 10% serum) was added and incubated overnight at 4°C. After washing, coverslips were incubated with secondary antibody, washed in PBS, then in water and ethanol. Coverslips were mounted with DAPI Fluromount (Southern Biotech, 0100–020).

#### Quantification of γH2AX foci, micronuclei and nuclear area

Quantification was performed by visual examination under Axio Zeiss Imager.M2 microscope. The number of γH2AX positive NSCs was analyzed by scoring at least 100 cells per line in five independent biological replicates. The number of cells containing micronuclei was analyzed in at least 600 cells per line in three independent replicates. Nuclear area was calculated using a macro and scored in at least 100 cells per line in three independent replicates.

#### Confocal imaging of mitotic errors

For acetyl-α-tubulin and phospho-histone 3 immunostaining, cells were seeded onto coverslips in a 6 well plate. The coverslips were then fixed for 20 minutes with 4% PFA. Next, a blocking buffer (1x PBS, 5% normal goat serum, 0.3% Triton X-100) was prepared and added to the coverslips for 1 hour at room temperature. The blocking buffer was then removed and anti-Acetyl-α-Tubulin and anti-Phospho-Histone 3 primary antibodies were diluted to 1:400 and 1:200 respectively in antibody dilution buffer (1x PBS, 1% BSA, 0.3% Triton X-100) and both were added simultaneously to each coverslip. Coverslips were incubated with the primary antibody overnight at 4°C. Then cover slips were washed thrice for 5 minutes in 1X PBS. Subsequently goat anti-mouse and anti-rabbit secondary antibodies were diluted to 1:500 in the antibody dilution buffer and both were added simultaneously to each coverslip. Coverslips were incubated for 2 hours in the dark at room temperature with the secondary antibody and were then washed thrice for 5 minutes in 1X PBS. Then they were rinsed in double distilled H2O followed by 100% ethanol and left to air dry. Coverslips were then mounted onto microscope slides using DAPI fluoromount and left for 1 hour in the dark before imaging. Imaging was performed on an Axio Zeiss Imager.M2 microscope and on a Leica SP8 confocal microscope.

#### MTT assay

Metabolic activity was analyzed 48 hours after cell seeding (Thiazolyl Blue Tetrazolium Bromide, Sigma-Aldrich, M5655). Absorbance was measured at 560 nm using a microplate reader (Mithras LB 940, Berthold technologies). Values from the blank measurements were subtracted from the average based on six technical replicates. Five biological replicates were obtained.

### Computational Methods

#### Whole genome sequencing and variant calling

Whole-genome sequencing data and whole-exome sequencing data were processed by the DKFZ OTP pipeline ^35^. Briefly, this workflow is based on BWA-MEM (v0.7.15) for alignment, biobambam (https://github.com/gt1/biobambam) for sorting and sambamba for duplication marking. Copy number variants were called using ACESeq2 and structural variants were called by SvABA(v134) based on the aligned genomes. ACESeq2 output was used only for ShatterSeek and for all other analyses copy number variants were called using CNMops(1.32.0) ^36^ with a 20kb bin size, in combination with GC content correction and replication timing correction provided by ACESeq2. DNAcopy algorithm was used for copy number segmentation.

#### Inference of chromothripsis in bulk WGS data

Chromothripsis scoring of whole genome sequenced tumors were performed by ShatterSeek ^3^. Copy number variants from ACESeq2 ^37^ and structural variants from SvABA were provided as input to ShatterSeek. We applied the criteria from previous studies to define chromothripsis positive chromosomes ^38^ from ShatterSeek output.

#### Inferring double minute chromosomes by AmpliconArchitect

To infer double minute chromosomes by AmpliconArchitect, we provided AmpliconArchitect ^39^ genome segments with copy number >=3 and the tumor alignment as input. AmpliconArchitect was allowed to explore other genomic regions connecting to the candidate genomic segment in an attempt to construct a circular amplicon. The output from AmpliconArchitect was filtered by removing circular segments with average copy number less than 3 and non-circular segments.

#### Bulk RNA-seq Gene expression analysis

Bulk whole transcriptome sequencing data were processed and normalized by kallisto ^40^. GENCODE basic version 30 ^41^ along with Human genome reference GRCh38 were provided to kallisto as reference. Kallisto reported expression levels per transcript. The expression across transcripts were summed to produce a gene-level expression measurement in transcripts per million (TPM) and in raw count. We utilized DESeq2 for differential gene expression analysis ^42^. Genes were filtered if they did not exceed 10 counts in 3 or more samples to ensure good quality data. Chromothripsis positive medulloblastomas were compared to non-chromothriptic medulloblastomas in a multivariate analysis adjusting for tumor cell content as well as SHH subgroup (**Supplementary Table 4**). The top 2000 up- and down-regulated genes were subsequently used for GSEA enrichment analyses, equivalent to the single-cell data analysis described in *‘Differential gene expression analysis and Gene Set Enrichment Analysis for individual copy number clones’* (see below).

#### Copy number assessment of exome sequencing data

Control-FREEC version 11.4 ^43^ was used for copy number assessment of the high coverage exome sequencing data. Known SNPs from dbsnp v142 were used as reference. The analysis was restricted to the exome capture region without the untranslated regions (UTRs).

#### Phylogenetic inference from bulk sequencing data

Phylogenetic inference was based on SNVs and CNVs using Expectation-Maximization on a multinomial model as previously published in Körber et al., 2019 ^44^. A few adjustments were made to the inference algorithm, in order to account for multiple samples. These adjustments are outlined in the following and an updated version of the code is available on github (https://github.com/hoefer-lab/phy_clo_dy/multi_sample).

##### Input data

We used the read counts at all SNVs passing the quality filters for tree learning. If a mutation was absent in at least one sample, we manually checked whether the mutation was present in that sample but did not pass the filtering criteria and adjusted the input data accordingly. Coverage ratios were looked up at each mutated position using the output of Control-FREEC. In order to map copy number changes that did not carry an SNV to the tree, we additionally added all loci at which the coverage ratio changed in at least one sample to the input. If these positions did not harbor a mutation, we set the reference read counts to the average coverage across all mutated sites and the mutated read counts to zero. Moreover, we added the location of *TERT* and *PTEN*, as well as locations on chromosome 2p, 3p, 3q, 5p, 7p, 7q, 10p, 10q, 11p, 16q and 17p to map gains and losses on these chromosomal arms to the phylogenetic tree. The positions were taken as the midpoint of the gained or lost segment according to the output of Control-FREEC.

##### Candidate trees

The number of binary trees grows fast with the number of clades and thus finding the best solution in a multi-sample problem requires an efficient searching strategy. We here addressed this problem by initiating the algorithm with a set of candidate trees, which were based on prior information and sequentially expanded during the fitting procedure.

- A first set of candidate trees was based on the mutational spectrum across the three samples. Mutations were either shared by all samples, or by the primary tumor and the metastasis only, or were private to a single sample. Thus, the mutation spectrum is consistent with a phylogenetic tree setting the relapsed tumor apart from the primary tumor and the metastasis. This is the simplest tree that is in agreement with the data from a combinatorial point of view. In order to account for more complex solutions, we extended this basal tree to more complex candidate trees by splitting individual clades.

- Second, we took all unique trees consisting of up to five clades and split each clade into three subclones, corresponding to the three samples. These trees were extended by adding clades above each node during the optimization algorithm and accepted if they yielded an improved solution based on a Bayesian Information Criterion. Extensions were abrogated if each sample consisted of three subclones or if the solution did not improve.

##### Additional adjustments

As compared to the algorithm described in Körber et al., 2019 ^44^, we added a few additional adjustments:

- Model selection was based on a modified Bayesian information criterion as outlined in Körber et al., 2019, but without prior tree selection based on clonal mutation estimates.

- We accounted for the possibility that a mutation call was false negative in the candidate tree (i.e., truly present, but not detected in a sample).

- We restricted the range of normal copy numbers from [0.9, 1.1] to [0.95, 1.05].

- We added prior information on whether a copy number change observed in multiple samples was likely due to a single event based on manual inspection of the copy number profiles. Specifically, we required that the losses on 2p, 3p, 11p, 16q and 17p, as well as the gains on 3q were due to single events.

#### DNA methylation

The majority of the DNA methylation profiles were published in a previous study ^5^. Genomic DNA was extracted from fresh-frozen or formalin-fixed and paraffin-embedded (FFPE) tissue samples. DNA methylation profiling of all samples was performed using the Infinium MethylationEPIC (850k) BeadChip (Illumina, San Diego, CA, USA) or Infinium HumanMethylation450 (450k) BeadChip array (Illumina). All computational analyses were performed in R version 3.5.3 (R Development Core Team, 2021; https://www.R-project.org). Raw signal intensities were obtained from IDAT-files using the minfi Bioconductor package version 1.21.4 ^45^. Illumina EPIC samples and 450k samples were merged to a combined data set by selecting the intersection of probes present on both arrays (combineArrays function, minfi). Raw methylation signals were normalized by the function preprocessIllumina. Possible Batch-effects caused by the type of material tissue (FFPE/frozen) and array type (450k/EPIC) were adjusted by fitting univariable, linear models to the log2-transformed intensity values (removeBatchEffect function, limma package version 3.30.11). The methylated and unmethylated signals were corrected individually. Beta-values were calculated from the back-transformed intensities using an offset of 100 (as recommended by Illumina). Filtering of CpG probes was performed as described in Capper et al. 2018 ^46^. In total, 428,230 probes were kept for downstream analysis. To perform unsupervised non-linear dimension reduction, PCA was applied to the 50,000 probes with highest standard deviation and the resulting first 100 PCs were used for UMAP analysis (R package uwot 0.1.8). The following non-default parameters were applied: n_neighbors= 10; min_dist= 0.5.

#### Quality Control and scDNA-seq data processing

The raw base call files from the 10X Chromium sequencer were processed utilizing the Cell Ranger DNA (version 1.1.0) pipeline. First, the “cellranger mkfastq” command was used to demultiplex the sequencing samples and to convert barcode and read data to FASTQ files. Further, the “cellranger-dna cnv” command takes as input FASTQ files and performs reference alignment, cell calling, and copy number estimation. As a reference genome we used pre-build Human reference GRCh37 (hg19), which was downloaded from 10X genomics website (version 1.0.0 from June 29 2018, https://support.10xgenomics.com/single-cell-dna/software/downloads/latest). To improve the sensitivity of the CNV analysis, we modified the very stringent 10X Cell Ranger DNA filtering pipeline for the detection of noisy cells and defined our own set of quality control metrics. A cell was labelled as noisy if any of the below-mentioned criteria were met:

1. The number of breakpoints was larger than 2 standard deviations from the median number of breakpoints (on a log scale).
2. The “is_high_dimapd” value provided by the Cell Ranger DNA pipeline (in the per cell summary file) was set to true, and the number of breakpoints was larger than 1 standard deviation from the median number of breakpoints (on a log scale).
3. The “ploidy_confidence” value provided by the Cell Ranger DNA pipeline (in per cell summary file) was between 0 and 2, and the number of breakpoints was larger than 1 standard deviation from the median number of breakpoints (on a log scale).
4. The median ploidy of the cell was 0.
5. 20% of the original ploidy estimates were corrected using the procedure described below.

Ploidy estimates for sample LFSMBP-Nuclei both before and after the removal of noisy cells are shown in **Supplementary Fig. 1**.

Additionally, we further reduced likely technical variation by correcting ploidy values in bins with rare ploidy profiles (**Supplementary Fig. 1e**). First, we grouped high quality cells by their average ploidy (e.g. diploid group of cells, tetraploid group of cells, etc.). If normal cells were present (based on manual curation), they were always placed in a separate group. Second, we counted ploidy values for every bin coordinate (20Kb windows) across all cells within a group. If some ploidy value was present in less than 4 cells, we corrected it to the most common ploidy value for that genomic coordinate and group of cells. Only groups with more than 3 cells were considered for correction. Additionally, if no ploidy value was shared by 4 or more cells from a group, no correction was performed.

#### Clonal inference from scDNA-seq data and lineage tree reconstruction

To infer CNV clones, we started with normalizing cells to the same ploidy. Specifically, we normalized cells to the most common ploidy among cells from a sample, which in our case was either two or four, depending on the sample (c.f. **Supplementary Table 1**).

To normalize cells to diploid, we applied the following formula:

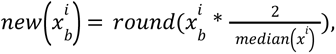

for cell *i* = 1, …, *n* and bin *b* = 1, …,*m*. Here, *x* indicates the ploidy value for the bin *b* in cell *i*.

Similarly, we normalized cells to tetraploid by applying

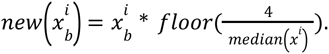

The distance between cells was estimated as the fraction of bins that are different between two cells:

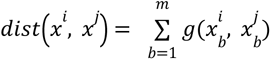

where 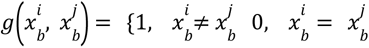

We then performed complete-linkage hierarchical clustering with the above defined distance measure, using the hclust function from R (4.0.0). We set a threshold *d* on the maximum permitted distance between cells in the same cluster to cut the dendrogram, and we defined clones as clusters with more than *k* cells. All cells that are assigned to clusters with less than *k* cells were discarded as noise. Thresholds *d* and *k* were defined ad-hoc with the objective of maximising the number of kept cells in the clones while at the same time capturing clonal diversity. A lower *d* value will generate more clusters but also increase the number of small clusters that will be discarded. A lower *k* value will generate smaller clones that might only capture noise. This approach was inspired by CHISEL ^47^. The selected *d* and *k* values for all samples are shown in **Supplementary Table 1**.

Next, for each clone we assigned a consensus CNV profile by taking the most common ploidy from all cells in that clone, for each bin. If a clone consisted of only two cells, the CNV profile of the cell with the smaller number of breakpoints was chosen as the clone CNV profile. If an unmappable region was found between two CNV events of the same ploidy, we estimated its ploidy to be the same as the surrounding CNV events (e.g. 2-2-4-**4**-NA-NA-**4**-4-4-2-2 where NA is an unmappable region, would be corrected to 2-2-4-4-**4**-(4)-(4)-**4**-4-4-2-2). To calculate the distance between any two clones we first intersected the clone’s CNV profiles to define a shared feature space (**Supplementary Fig. 1f)**. For this we used intersectBed from bedtools (version 2.24.0). We noticed that there is an inflation of short genomic regions after intersection, which is most likely caused by imprecise breakpoint detection. Depending on the sample-specific distribution of genomic region lengths we set a minimum length threshold to filter out genomic regions (see **Supplementary Table 1**). We then calculated the distance between two clones *C^α^* and *C^β^*as

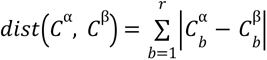

where *r* corresponds to the number of genomic regions after discarding short regions and 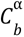 indicates the ploidy value for clone *α* and genomic region *b*.

Finally, we fitted a minimum spanning tree (MST) over clones using the above defined distance. We added one artificial zygote clone that is forced to become the root of the tree. Initially, the MST was constructed using only diploid clones (where the majority of the cells were diploid before normalization). Subsequently, tetraploid clones (with majority of the cells being tetraploid before normalization) were attached to their closest clone in the tree.

#### Inter LFSMB samples clonal matching

For every pair of LFSMB samples (including primary, relapse, patient tumor and PDX) we built a lineage tree combining clones from both datasets, following the algorithm described in the previous section. Based on the distribution of pairwise distances intra and inter clones and their CNV heatmaps (**Supplementary Fig. 2a**) we performed a visual inspection to select the clones that were the closest based on shared CNVs and assigned them as matching in Fig 1c. In the case of C4 in LFSMB_P_PDX we were able to match it best to a subclone within C5 in LFSMB_P_Nuclei.

#### Chromothripsis (CT) detection at the single-cell and clone levels

We adjusted established criteria for inferring chromothripsis in cancer genomes from bulk whole-genome sequencing ^6–8^. scDNA CNV data consists of stretches of CNV values per bins (20kb) across the genome, divided into chromosomes. As so, every cell is represented as a stretch of discrete values that represent the CNV states that are identified in all bins, at every chromosome. To detect CT in a cell we assess how many CNV state changes are detected within a 50Mb window. As an example, the following stretch “22222233232222” contains 5 switches. If 10 switches are observed within a 50Mb window that window is considered to be chromothriptic. We implement a sliding window approach where we assess every possible 50MB window within every chromosome and assees in each case whether or not it is chromothriptic. We finally compute a CT chromosome score calculated as the fraction of evaluated windows in a given chromosome that are positive to CT. Note that this score denotes how chromothriptic a chromosome is (Fig 1e-f).

In addition to the chromosome level CT score we have also implemented a bin level score (Fig 1d). The methodology is the following: When sliding a window to score CT at the chromosome level every bin is evaluated multiple times as many windows will contain it (n_windows=window_size / bin_size). We calculate the bin CT score as the fraction of windows containing a single bin that are CT positive following the aforementioned criteria. This score is assigned to every bin but represents the level of CT in a window_size neighborhood, centered in the bin of interest. Note also that there is a border effect due to bins in the limits of the chromosome being evaluated less times and hence CT can be underestimated in them. The bins affected by this effect will be the ones located in the regions chr_start / chr_start+window_size and chr_end-window_size / chr_end.

Given that single cell data can be quite noisy we implement two strategies to have a better signal to noise ratio.

Strategy 1: The first strategy is to filter out switches that are too short (i.e only within a few bins) which might be an indication of a noisy signal. We introduce the parameter *NConsecutiveBins* to define how many bins a given state needs to persist in order to be considered as valid. If we take the same example from before, “22222233232222” with *NConsecutiveBins*=1 we observe 5 switches whereas with *NConsecutiveBins*=2 we observe 3 switches. This is easily implemented by first compressing the CNV stretch from “22222233232222” to “2**6**3**2**2**1**3**1**2**4**” where when taken in pairs, the first number denotes the CNV state and the second the number of times it is repeated. We can then count how many pairs are left after imposing to keep only those such as their repeat number >= *NConsecutiveBins.* STP-Samples were scored using NConsecutiveBins=1. MB243-Nuclei and RCMB18-PDX were scored using NConsecutiveBins=30.

Strategy 2: The second strategy consists in aggregating groups of cells that are similar to each other (e.g the ones belonging to the same clone) and create a consensus CNV profile for them by majority vote (i.e for the aggregated profile we keep the CNV state that has the majoritarian representation at each bin).

#### Subsampling to estimate the robustness of the chromothripsis scoring

After grouping cells into clones, we calculated CT scores at the chromosome level by aggregating CNV profiles of cells belonging to the same clone by applying a majority vote strategy (Fig 1e). To evaluate how robust our CT scoring is, we took all clones in LFSMB_P_Nuclei and subsampled cells within them without replacement to create new clones containing 80% of the original cells. We iterated the process 100 times. In every iteration the CNV profiles from the subsampled cells were aggregated into majority vote CNV profiles and scored for CT. The median CT score values for the subsampled clones, across 100 iterations, were highly correlated with the original CT scores from the full dataset (**Supplementary Fig 2b**), indicating that the CT scoring strategy is robust.

#### Quality Control and scRNA-seq data pre-processing

The raw base call files from the 10X Chromium sequencer were processed utilizing the Cell Ranger Single-Cell Software Suite (release v3.0, https://support.10xgenomics.com/single-cell-gene-expression). First, the “cellranger mkfastq” command was used to demultiplex the sequencing samples and to convert barcode and read data to fastq files. Based on the fastq files, “cellranger count” was executed to perform alignment, filtering, as well as barcode and unique molecular identifier (UMI) counting. The reads from single-nuclei RNA-sequencing were aligned to the pre-mRNA hg19 reference genome, while the reads from single-cell RNA-sequencing were aligned to the hg19 reference genome, implementing a pre-built annotation package downloaded from the 10X Genomics website. For all single-cell RNA-sequencing data resulting from PDX samples, we also mapped the reads to the mouse genome (mm10) in order to check whether cells map better to human or mouse. If less than 1% of the reads were aligned to hg19, we defined the respective cells as mouse cells. Several output files including a barcoded binary alignment map (bam) file and a summary csv file are provided. Importantly, a filtered feature-barcode matrix folder, containing a valid barcode file for all QC-passing cells, a feature file with ensembl gene ids and a matrix in the genes x cellsformat are generated. The filtered genes x cells matrix was further used as input for the data processing workflow described in the following.

#### Analysis of single sample scRNA-seq data using scanpy

The output from the Cell Ranger analysis framework was used as input to a custom analysis workflow, structured around the scanpy software toolkit in python ^48^. First, genes that were expressed (>=1 count) in <=5 cells across the whole dataset were removed (sc.pp.filter_genes with min_cells=5). Next, we filtered single-cells and single-nuclei data individually. For single-nuclei, we filtered them for i) counts (500 < total_counts < 25,000), ii) genes (300 < n_genes < 6000), iii) mitochondrial genes (pct_counts_mt < 5%) and ribosomal genes (pct_counts_ribo < 10%). In contrast, single-cells were filtered differently, but also for i) counts (500 < total_counts < 25,000), ii) genes (200 < n_genes), iii) mitochondrial genes (pct_counts_mt < 10%) and ribosomal genes (pct_counts_ribo < 40%). In addition, we used scrublet ^49^ to remove potential doublets in our dataset, see **Supplementary Table 3** for details).To account for variable sequencing depth across cells, we normalized unique molecular identifier (UMI) counts by the total number of counts per cell, scaled to counts per 10,000 (CP10K; sc.pp.normalise_per_cell), and log-transformed the CP10K expression matrix (ln[CP10K+1]; sc.pp.log1p). Next and to generate cell type clusters, we selected the 2,000 most variable genes across samples by (1) calculating the most variable genes per sample and (2) selecting the 2,000 genes that occurred most often across samples (sc.pp.highly_variable_genes). After mean centering and scaling the ln[CP10K+1] expression matrix to unit variance, principal component analysis (PCA; sc.tl.pca) was performed using the 2,000 most variable genes. To select the number of PCs for subsequent analyses, we used a scree plot and estimated the “knee/elbow” derived from the variance explained by each PC. Finally, we used bbknn v1.3.9 ^50^ to batch align samples and control for sample specific batch effects, following the recommendations of Luecken et al. ^51^. Following this, we calculated clusters using the Leiden graph-based clustering algorithm v0.7.0 ^52^, which were subsequently used for differential gene expression as described in the following.

#### Differential gene expression analysis and cell cluster annotation

To evaluate the cellular identity of distinct clusters, we annotated them based on the expression of known cell marker genes collected from the literature. For this purpose, we performed a two-sided Wilcoxon rank-sum (Mann-Whitney-U, Benjamini-Hochberg adjusted) test to compare each individual cluster to all other cells. Next, we then used the mentioned list of genes to assign cell identities to specific clusters. This list of known marker genes included *CD74, SAT1, MERTK* (macrophages), *VWF, EGFL7, INSR* (endothelial cells), *COL4A1, FN1, CDH11* (meninge cells), *FABP7, CLU, GFAP, SLC1A3, PTN* (astrocytes), *CD74, HLA-DRB1, RGS1, LYZ, CD81* (microglia), *GLI2, PTCH2, HHIP, POU6F2* (malignant SHH), *MKI67, TOP2A, DIAPH3, POLQ* (malignant cycling), *PTPRD, MARCH1, NCAM2, PLCL1, NEUROD1* (malignant neuronal development I) and *RASGEF1B, SLC26A3, LINGO1* (malignant neuronal development II). The expression of these marker genes was visualized using scanpy’s “DotPlot” function. In cases where the identity could not be resolved, the highest variable genes were used as input for a CellMarker database search (http://biocc.hrbmu.edu.cn/CellMarker/ ^53^). Alternatively, the ToppGene suite (https://toppgene.cchmc.org/) was used to evaluate the cellular identity of a cluster ^54^. As these databases comprise a collection of marker genes from distinct scRNA-seq studies, the cell type of a cluster could usually be assigned successfully.

#### Bulk chromothripsis signature analysis in single-cells

Using the genes present in the chromothripsis signature from bulk RNA-sequencing (see above), we investigated whether we could detect evidence for the aggressiveness of these tumors in our scRNA-seq data. For this purpose, we distinguished between up- and down-regulated genes by splitting them into a chromothripsis score [Upregulated] and [Downregulated]. Both lists of genes were intersected with the genes present in the scRNA-seq matrices for each sample. Following this, we utilized scanpy’s ‘scanpy.tl.score_genes’ function and the raw expression values to calculate the average expression of the chromothripsis associated genes subtracted by the average expression of a set of random reference genes. This generates a single-cell chromothripsis score, which we visualized on the UMAP embedding for the up- and down-regulated genes respectively.

#### Single-cell RNA-seq copy number detection by inferCNV

For the inference of copy number variation in single-cell RNA-seq data we used inferCNV, a method which uses variability in relative gene expression to assign copy number profiles to each cell ^27^. Following the vignette provided by inferCNV’s author (https://github.com/broadinstitute/inferCNV/wiki), the count matrix from the quality-controlled ScanPy objects, generated for each single sample, was used as input. In general, inferCNV is able to detect large-scale alterations, for instance amplifications or deletions of entire chromosome arms by comparing the gene expression of the cells of interest relative to a population of reference cells. In case of the single-nuclei data investigated here, we leveraged endothelial cells from the matching donor as a reference cell population, as they were present in each donor and presumably had no copy number variants (CNVs). In contrast, for the single-cell data from PDX samples, we used cells which were assigned as mouse cells. Following this further, inferCNV routinely filters low quality genes, normalizes for sequencing depth and applies a log-transformation (see https://github.com/broadinstitute/inferCNV/wiki for further details). Each cell is smoothed using a weighted running mean of 151 genes with a pyramidal weighting scheme, favoring genes according to their proximity, and finally centered according to the cell’s median expression. This assumes that most genes are not in CN altered regions. To avoid considerable impact of any particular gene on the moving average, the gene expression values were limited to [−3,3], hence replacing all values above 3 by 3, and all values below −3 by −3. Afterwards, CNVs were estimated based on the average normalized gene expression of the cells of interest relative to the normalized gene expression of the reference cells, referring to a subtraction of the mean of the normal from the tumor cells. Building up on this, dynamic noise filtering was applied to enhance the signal-to-noise ratio between potential CN altered regions and diploid regions. In addition, we leveraged the capability of inferCNV to use a hidden-markov model (HMM) and Bayesian Network to calculate the underlying probabilities for each determined CNVs. For each of the mentioned steps, including the modified expression, HMM and BayesNet, heatmap visualizations were generated by default. In addition to the dynamic de-noising, an additional median filter was used to adapt the visual representation of the modified expression. The output from this median filtering step was utilized as input to a custom plotting script in R.

#### Integrating scDNA- and scRNA-seq data based on copy number profiles

In order to integrate and project our scDNA copy number clones to the scRNA-seq data, we designed an approach which relies on the correlation of a cell’s individual scRNA-seq gene profiles to a clone scDNA-seq gene profile. In detail, we first transformed the scDNA-seq information from segment/bin to gene-level copy number, as genes are the common denominator between both sequencing outputs. In order to accomplish this, the gene-level coordinates for hg19 were used from biomaRt and transformed them into a GRange object using the GenomicRanges package in R. Similarly, the segment coordinates from the constructed segment x cells matrix were converted to a GRange object as well. Using the “mergeByOverlaps” function of the GenomicRanges package, both information layers were combined by mapping the respective segment to the matching genes. Then, the mean copy number value per gene was calculated based on the mapped segments. In the next step, we used the evaluated clones from **Figure 1**, to create an average copy number profile per gene for each clone, represented by a vector of clone x genes. Importantly, in samples with clones depicting a median ploidy equal 4, we normalized the ploidy to 2 as whole genome doubled populations can effectively not be detected in relative copy number inference (although it might be theoretically possible). In parallel, we prepared the modified gene expression results from inferCNV, a genes x cells matrix with continuous expression values between 0.9 and 1.1, for the correlation approach by discretizing it. Thereby, we assume that the relative copy number inference can only detect losses and gains. Hence, we assigned a diploid copy number value of 2, if the modified gene expression value was 1.0 - sd(modified_expression) ≤ modified_expression ≤ 1.0 + sd(modified_expression). In contrast, if the modified gene expression was smaller or greater than the respective boundaries, a copy number value of 1 and 3 was assigned respectively.

Given that many chromosomes in both data modalities will not be informative to differentiate clones, we subset the number of chromosomes for the correlation to a minimum. For this purpose, we assessed the variable chromosomes in the scDNA data. First, we related genes to the respective chromosome arms they are located on using coordinates from hg19 and the identical features as described for the segment transformation shown above. Next, we counted the number of distinct copy number values per chromosome arm and clone, resulting in a table of clones x copy number values. Using this table, we calculated the frequency of CNVs per clone per chromosome, which was subsequently utilized to assess the standard deviation in the CNV distribution across clones. If a chromosome of interest showed a variability greater than 15%, we included it in our correlation approach, while all other chromosomes were discarded. It should be noted that this threshold was applied to all samples, except from RCMB18-PDX (threshold = 1%). Lastly, we compared the results from this method to the visual information content of each chromosome in scRNA-seq. Thereby, if a chromosome was not informative, i.e. did not contain CNVs or did not display heterogeneous CNVs, the respective chromosome was excluded from the downstream correlation.

Using the subset of most variable genes, mapped to the variable chromosomes, we calculated the Pearson correlation between each copy number profile from the scDNA-seq clones and each cell in the scRNA-seq data (R_o_). In equivalence, and to compare the observed data to expected background signal, we leveraged the scDNA-seq data to create random scDNA clones. For this purpose, the real data clones were permuted to create 1000 newly derived copy number clones. Equivalent to the observed clones, we calculated the Pearson correlation between each permuted copy number profile and each cell in the scRNA-seq data (R_p_). Next, for each cell from scRNA-seq, the maximum Pearson correlation to one specific observed clone and simulated clone was utilized to create an empirical (observed) and null (permutated) distribution (Fig. 7a). Following this, we calculated a p-value for each clone-cell pair comparing the empirical and null distribution:

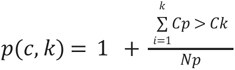

where p(c,k) is the p-value for a respective cell c correlated to a clone k, C_p_ is the correlation between the cell c and a permuted clone k_p_, C_k_ is the correlation between the cell c the observed clone k, while N_p_ is the total number of permutations. Lastly, each cell was assigned to its respective scDNA clone counterpart according to the maximum correlation and if the bonferroni corrected p-value (p(c,k) * number_clones) was smaller than 0.05. These assignments were then used to generate custom copy number heatmaps with the original modified gene expression values from inferCNV using a custom R script.

#### Differential gene expression analysis and Gene Set Enrichment Analysis for individual copy number clones

Using the copy number clone information generated as described above, we performed differential gene expression analysis between each individual copy number clone and all other cells. The two-sided Wilcoxon rank-sum (Mann-Whitney-U, Benjamini-Hochberg adjusted) test from scanpy was utilized in order to identify significantly up-regulated genes. For individual copy number clones with at least one significantly differentially expressed gene, the complete gene list was combined with the logFC values and utilized as input for gene set enrichment analysis (GSEA) as implemented in the R package HTSanalyzeR2 (https://github.com/CityUHK-CompBio/HTSanalyzeR2, ^55^). Thereby, GSEA was performed using the MSigDB hallmark gene sets provided by Liberzon and colleagues ^56^. The results from GSEA were filtered according to the underlying p-value (FDR < 0.05, Kolmogorov–Smirnov statistic, Benjamini Hochberg adjusted). Hence, if a pathway was significantly altered (p-value < 0.05), it was kept in the analysis, while non-significant pathways were discarded (p-value > 0.05). Subsequently, the clones and altered pathways were visualized using a custom R script.

#### Evaluating druggable targets from scDNA-sequencing data

CNVs within bins corresponding to genes reported as potential druggable targets by Worst et al 2016 et al ^20^ were evaluated across clones to evaluate presence or absence of focal gains. We calculated the ratio between the average CNV value for the gene and its corresponding Chr CNV, for all genes across cells in our datasets as Gene2Chr_ratio=log2(GeneCNV / Chr CNV). This ratio allows us to select genes with focal gains. We kept those genes for which ratio >= 1 for at least 1 cell. For the genes that fulfilled the aforementioned criteria we calculated their Gene2Chr_ratio for all cells and grouped them by DNA clone. We observed that in some cases genes show clonal gains across clones whereas others show subclonal differences (Fig 1H).

#### Analyzing the expression of druggable targets from scDNA

We investigated the expression of druggable targets from scDNA-sequencing (see above) in the scRNA-seq data to evaluate whether we can observe transcriptional consequences. Hence, we utilized the normalized expression matrix from scRNA-seq for each sample and subset it to the druggable target genes. Then, we used the integrated clone as well as the cell type information for normal cells to compare the expression of each of these groups of cells. Importantly, we removed cells with no expression in a respective gene as well as genes which were not expressed in at least 50 cells. Lastly, we combined the expression and CNV visualizations using a custom R script.

#### Assessing the influence of copy number variation in double-minute chromosomes on gene expression

In order to assess the transcriptional consequences of double-minute (DM) chromosomes, we evenly analyzed the scDNA- and scRNA-seq data. We determined the genes which are potentially influenced by DM chromosomes by intersecting the DM coordinates in scDNA-seq with the gene coordinates from hg19. Following this, we calculate the mean CNV value for each gene in this list by averaging bins within the respective gene. Importantly, we calculated these values on a per-clone level. Next, we leveraged the average CNV values per gene per clone to compare them to mean normalized gene expression values from the respective integrated clone in scRNA-seq. Finally, we investigated a correlation between both gene expression and CNV in each sample respectively utilizing Pearson’s correlation.

## DATA AVAILABILITY

Sequencing data **will be available (data deposition in progress)** in the European Genome-phenome Archive (EGA), which is hosted by the EBI and the CRG, under accession number EGAS00001005410. All other data are available within the Article, Supplementary Information or available from the authors upon reasonable request.

## CODE AVAILABILITY

The aforementioned computational methods provide a summary of the procedures implemented in various custom-made R, python and bash scripts. These scripts contain the commands run for the analyses highlighted in this publication. In order to sustain reproducibility, they are publicly available on Github (https://github.com/PMBio/MB_scSeq).

## ACKNOWLEDGEMENTS

We thank Peter Lichter and Aurelio Teleman for discussions, Frauke Devens, Michaela Hergt, Brigitte Schoell and Katharina Bauer for technical support, the Sequencing and the Microarray units of the Genomics and Proteomics Core Facility (DKFZ), the EMBL Sequencing facility, the DKFZ FACS Core Facility and the DKFZ Imaging Facility. This research was supported by the DFG, the Wilhelm Sander Foundation and the Fritz Thyssen Foundation.

## SUPPLEMENTARY MATERIAL/FIGURES

Separate file

