## Supplementary Figures for "Single cell multi-omics analysis of chromothriptic medulloblastoma highlights genomic and transcriptomic consequences of genome instability"

### Supplementary Material

#### Supplementary Figures

##### Supplementary Figure 1

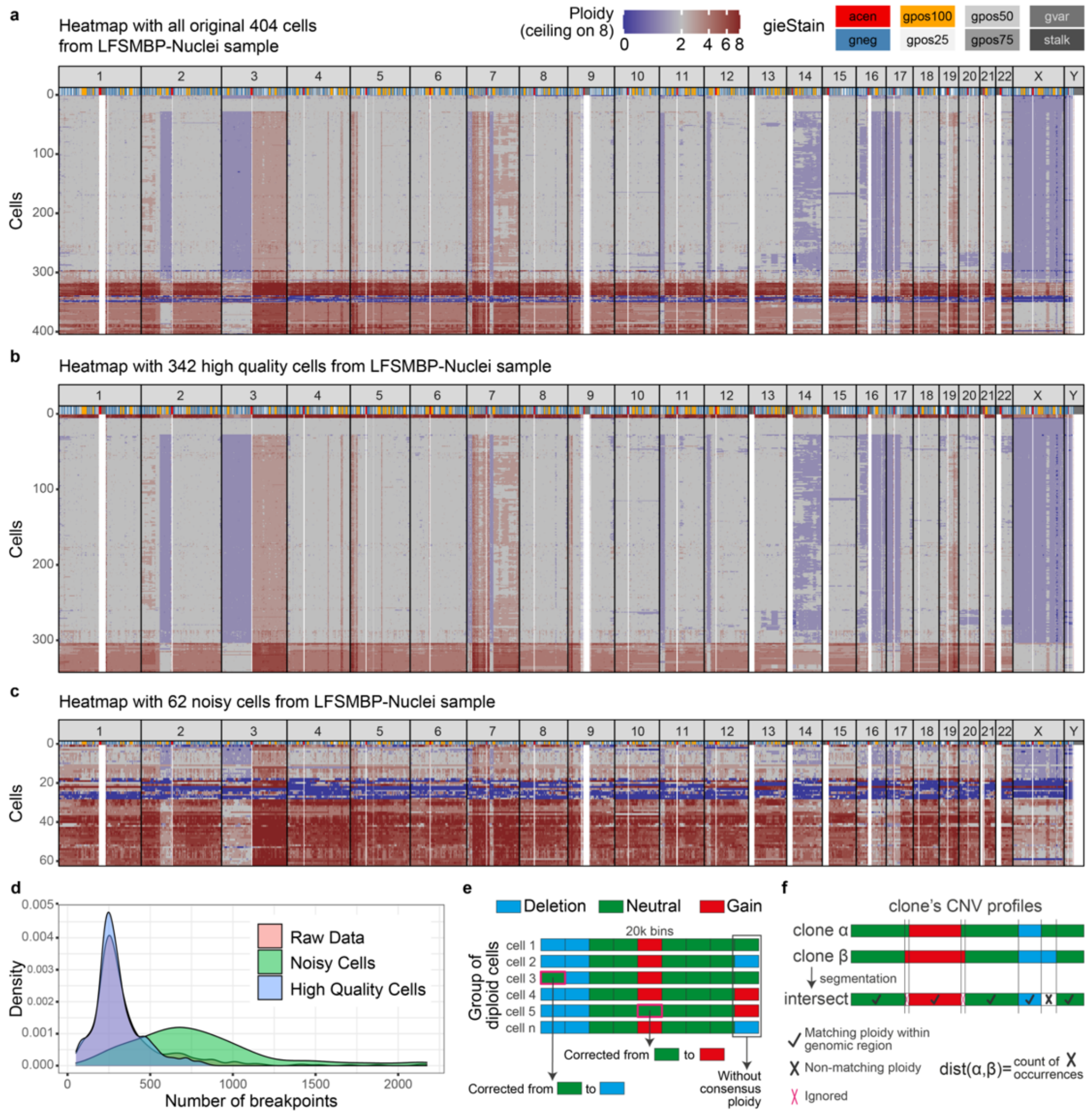

**Supplementary Figure 1. Single-cell data quality checks and filtering criteria. a,** Heatmap for all original 404 cells in LFS_MBP-Nuclei (before filtering). **b,** Heatmap for the 342 high quality cells in LFSMBP-Nuclei after quality check filtering. **c,** Heatmap for the 62 cells flagged as noisy in LFS_MBP-Nuclei. **d,** Distribution of breakpoints in the raw data as well as in the high-quality and noisy cell sets. **e,** Scheme of the process applied for correcting rare aberrant ploidies across bins in order to decrease noisy signal. The reason for doing this is that rare ploidies affect majority vote CNV profiles calculation for the clones and hence affect also the distances between clones (explained in panel f). Rare ploidy values, present in only a few cells (green example in cell 3 within the red rectangle in the first bin) are corrected to the most frequent value across cells, in order to decrease noise for clone matching. **f,** After rare ploidy correction, we calculate distances between all pairs of clones as a starting point for calculating clone lineage trees. We count the number of matching ploidies across the CNV majority vote vectors between two given clones (see the "**Quality Control and scDNA-seq data processing"** section for further details).

**Supplementary Figure 2.** Heatmaps (copy-number) and trees for all samples

**a  MB243-Nuclei**

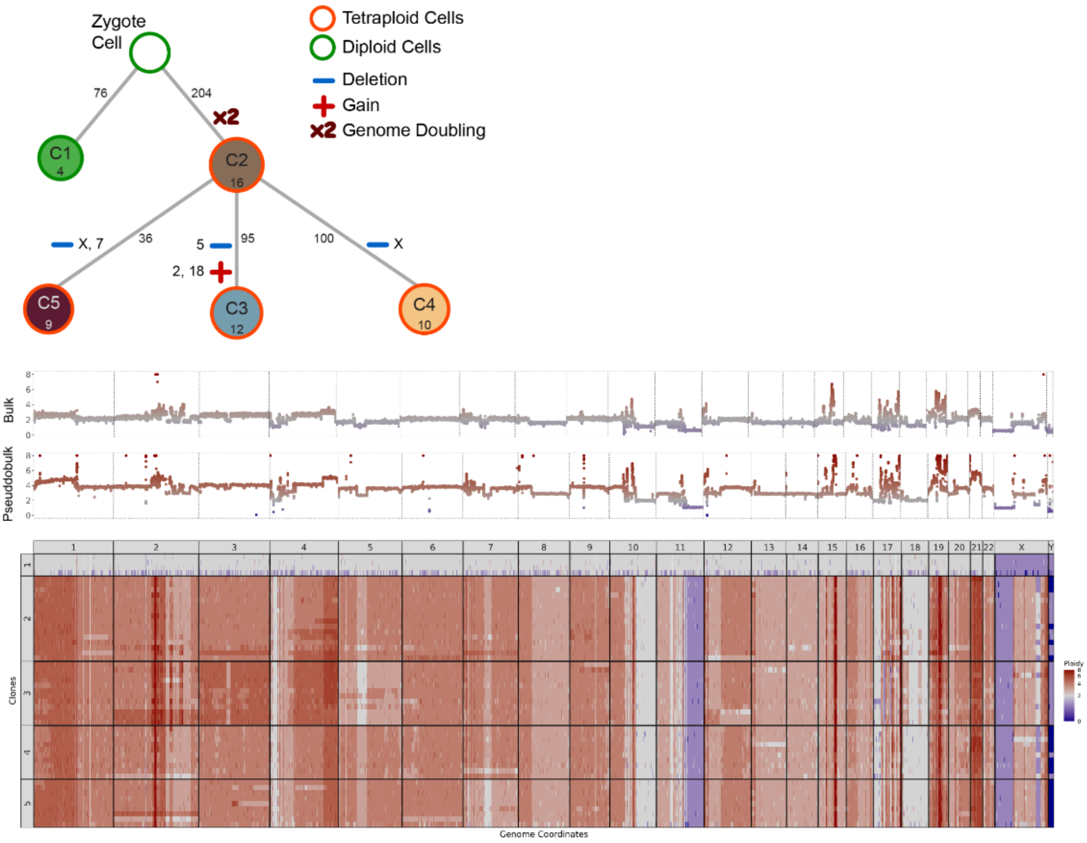


**b. LFSMBP-PDX**

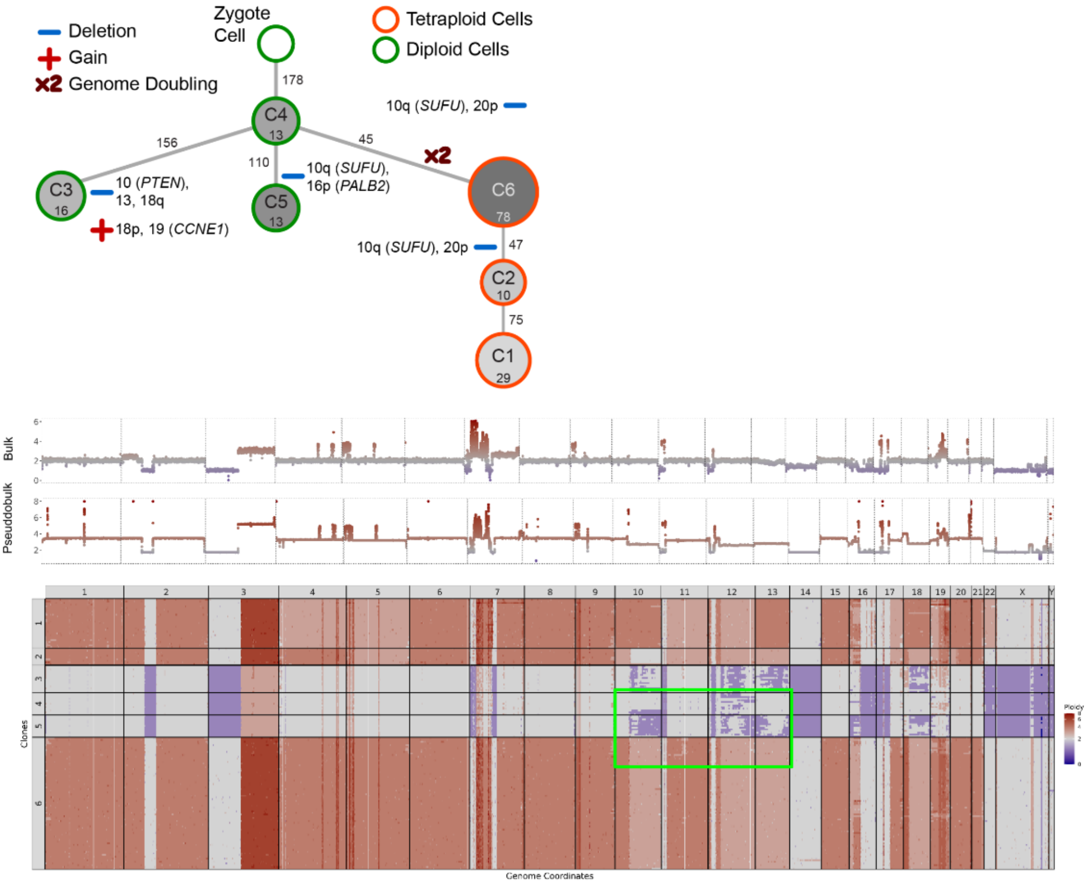

**Supplementary Figure 2.** Heatmaps (copy-number) and trees for all samples

#### c. LFSMB1R-Nuclei

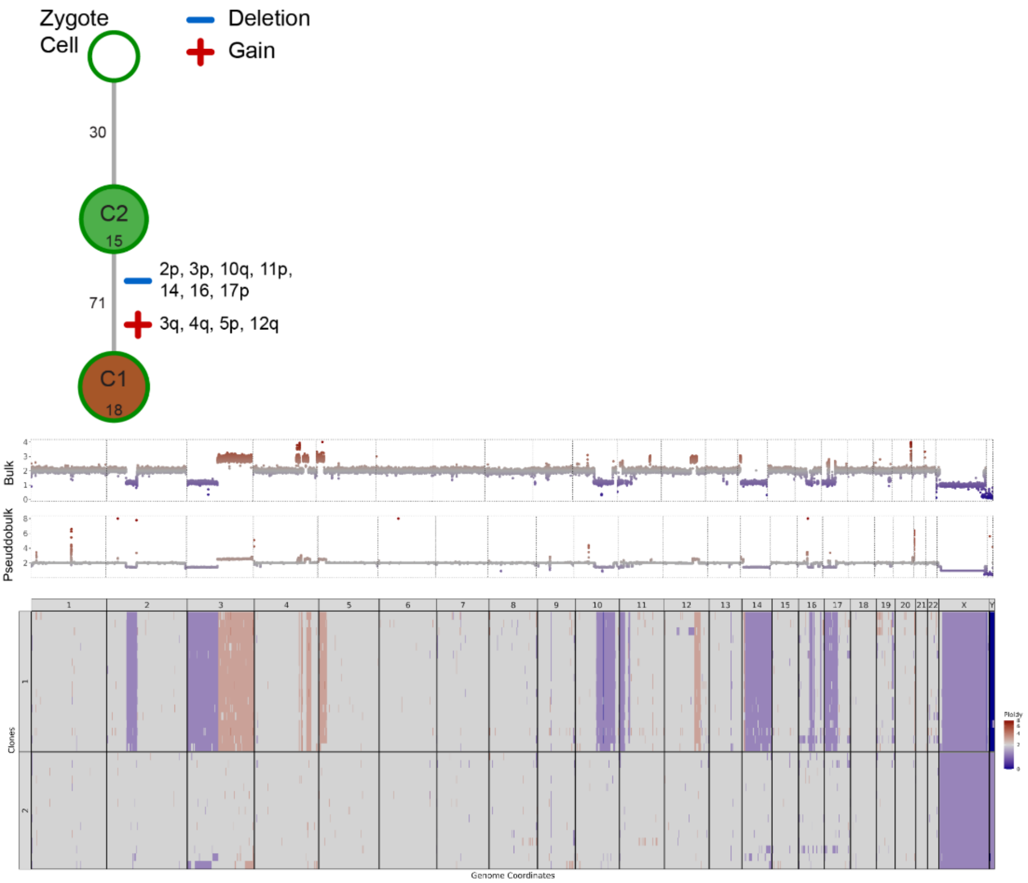


#### d. LFSMB1R-PDX

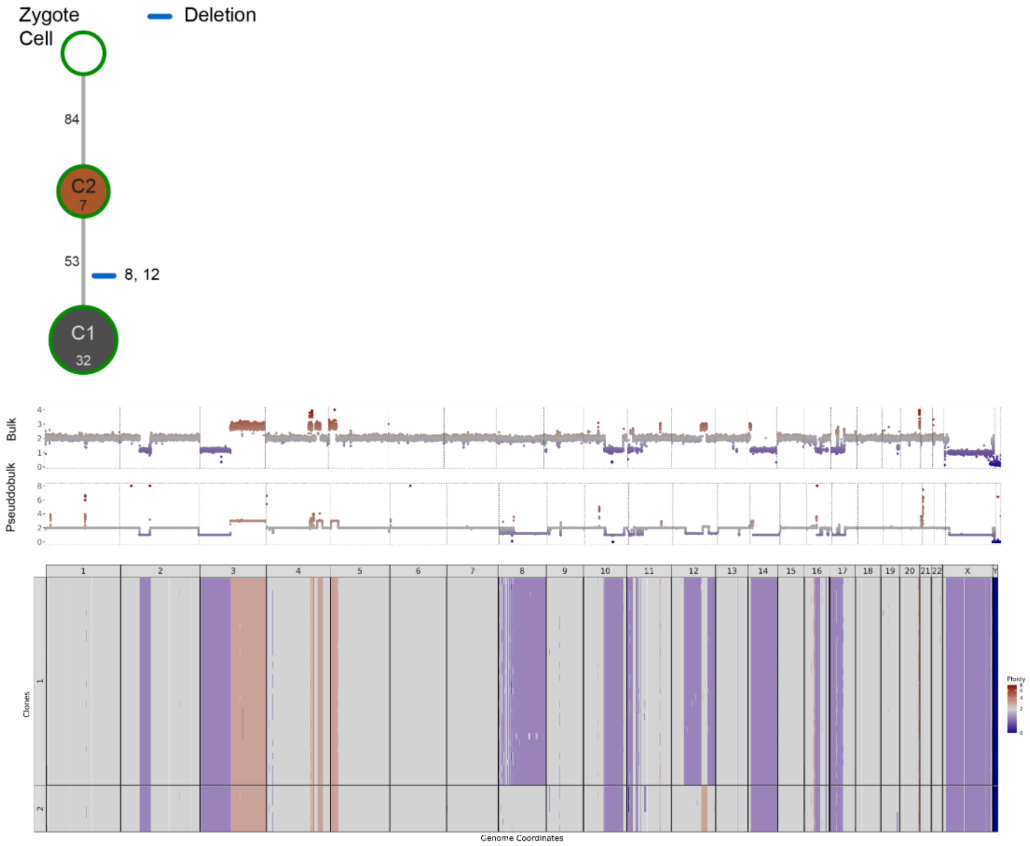

**Supplementary Figure 2.** Heatmaps (copy-number) and trees for all samples

#### e. RCMB18-PDX

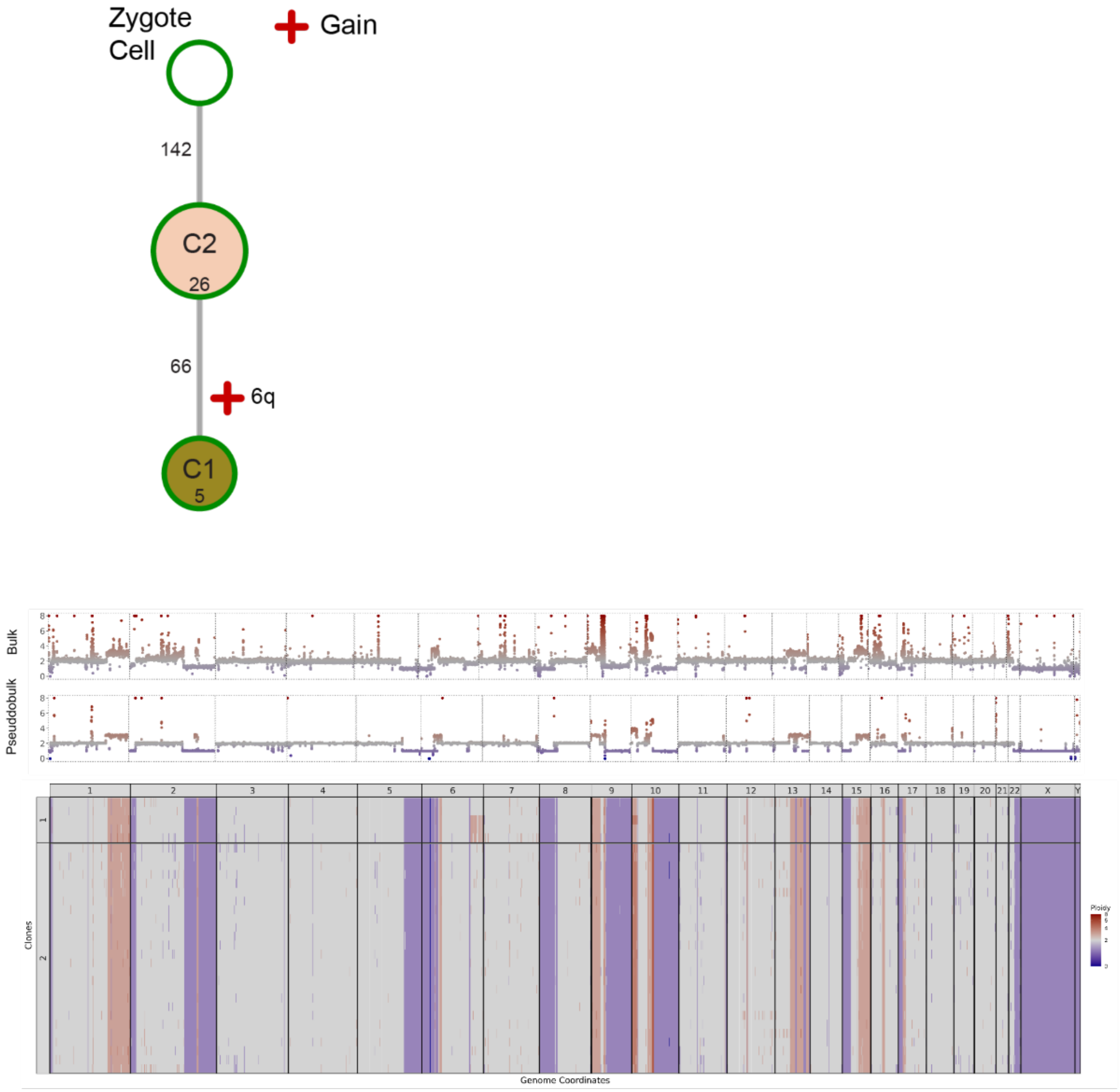


#### f. CT scoring in LFSMB_P_Nuclei subsamplings

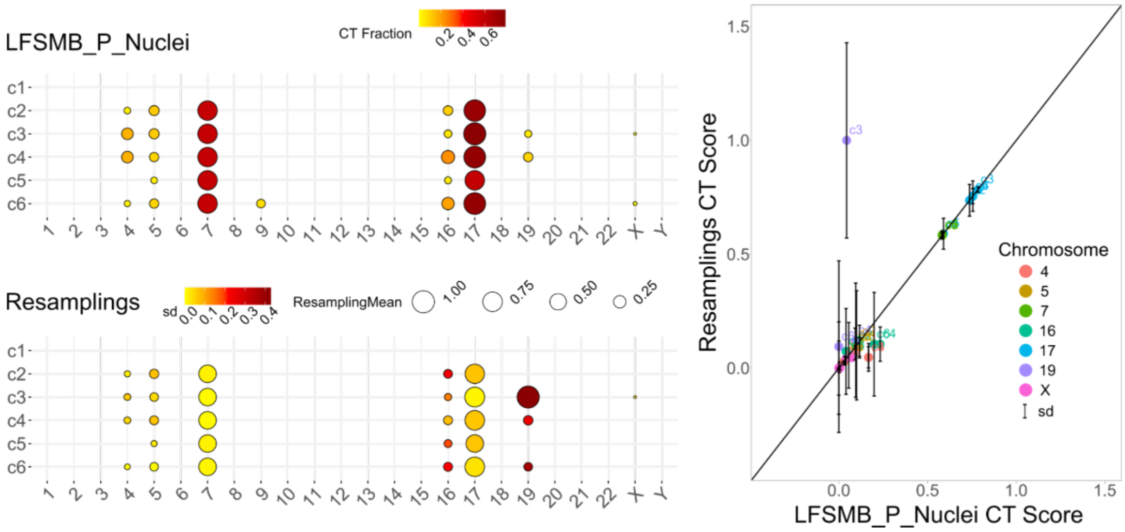

**Supplementary Figure 2. Copy number heatmaps and phylogenetic trees for all samples profiled by 10X scDNA. a,** Clonal tree structure and heatmap for all high-quality cells clustered by clone for all MB243-Nuclei. **b,** Clonal tree structure and heatmap for all high-quality cells clustered by clone for all LFS_MBP-PDX. **c,** Clonal tree structure and heatmap for all high-quality cells clustered by clone for all LFS_MB1R-Nuclei. **d,** Clonal tree structure and heatmap for all high-quality cells clustered by clone for all LFS_MB1R-PDX. **e,** Clonal tree structure and heatmap for all high-quality cells clustered by clone for all RCMB18-PDX. **f,** Comparison between CT Scoring for the LFS_MBP-Nuclei sample (up-left) and CT Scoring of 100 resamplings (subsampling of 80% of cells per clone without replacement, bottom-left). Right: scatterplot comparing chromosomes with non-zero CT values in LFS_MBP-Nuclei against the corresponding CT Score values.

#### Supplementary Figure 3

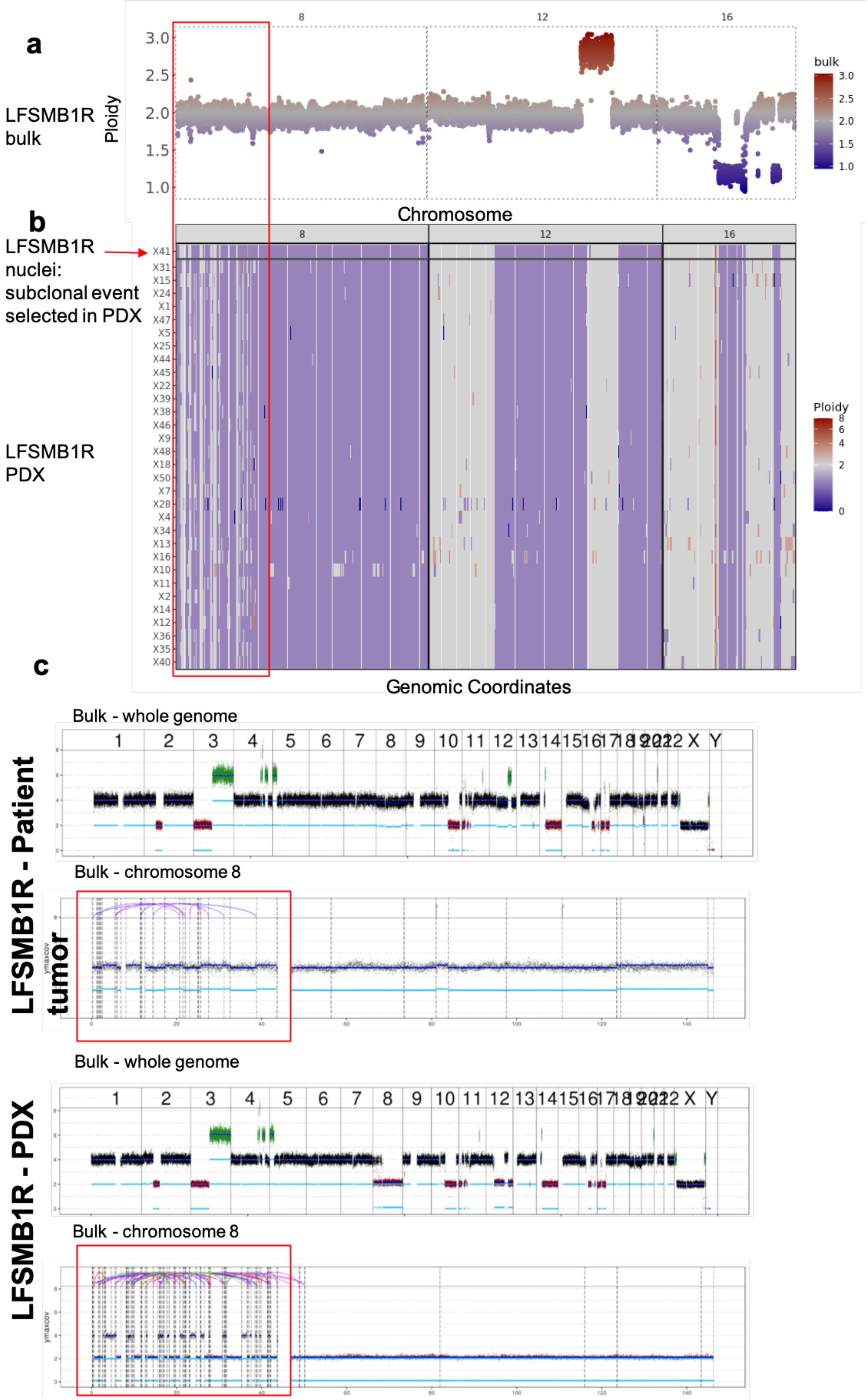

**Supplementary Figure 3. Validation of subclonal chromothriptic events in LFS_MB1R. a-b,** Representative example of subclonal chromothriptic (CT) event on chromosome 8 in LFS_MB1R-Nuclei sample. This CT event is not detected in bulk WGS data, which shows only minimal copy-number changes (bulk scatterplot, panel **a**). **c**, The subclonal event detected in LFSMB1R-Nuclei (heatmap, upper lane in **b**) is detected in the major clone in the matched PDX model (LFSMB1R-PDX), likely due to selection of the clone with chromothripsis on chromosome 8 in the xenograft.

**Supplementary Figure 4. Intra-tumor heterogeneity of druggable genes and double-minute chromosomes**

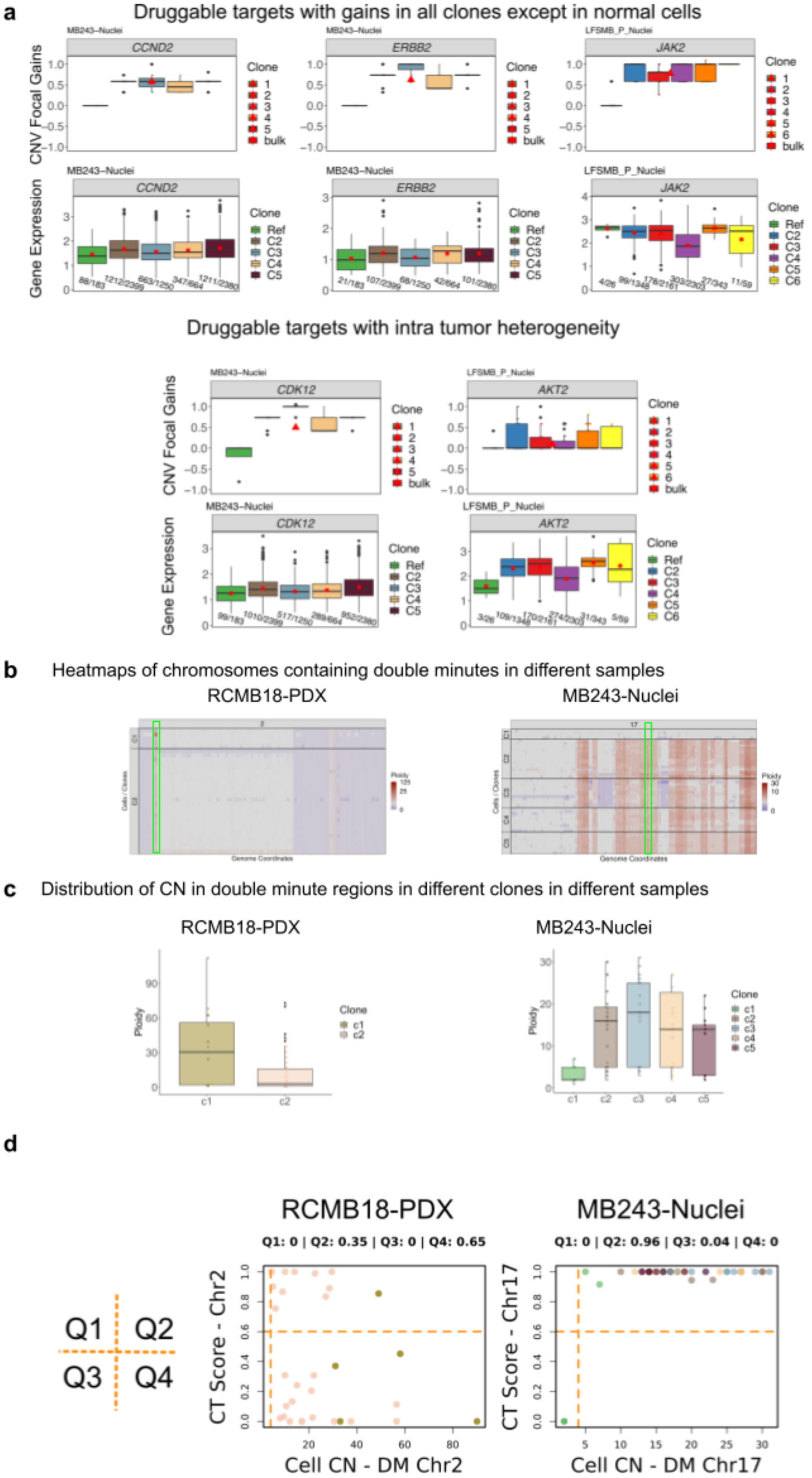

**Supplementary Figure 4. Intra-tumor heterogeneity of druggable genes and double-minute chromosomes**

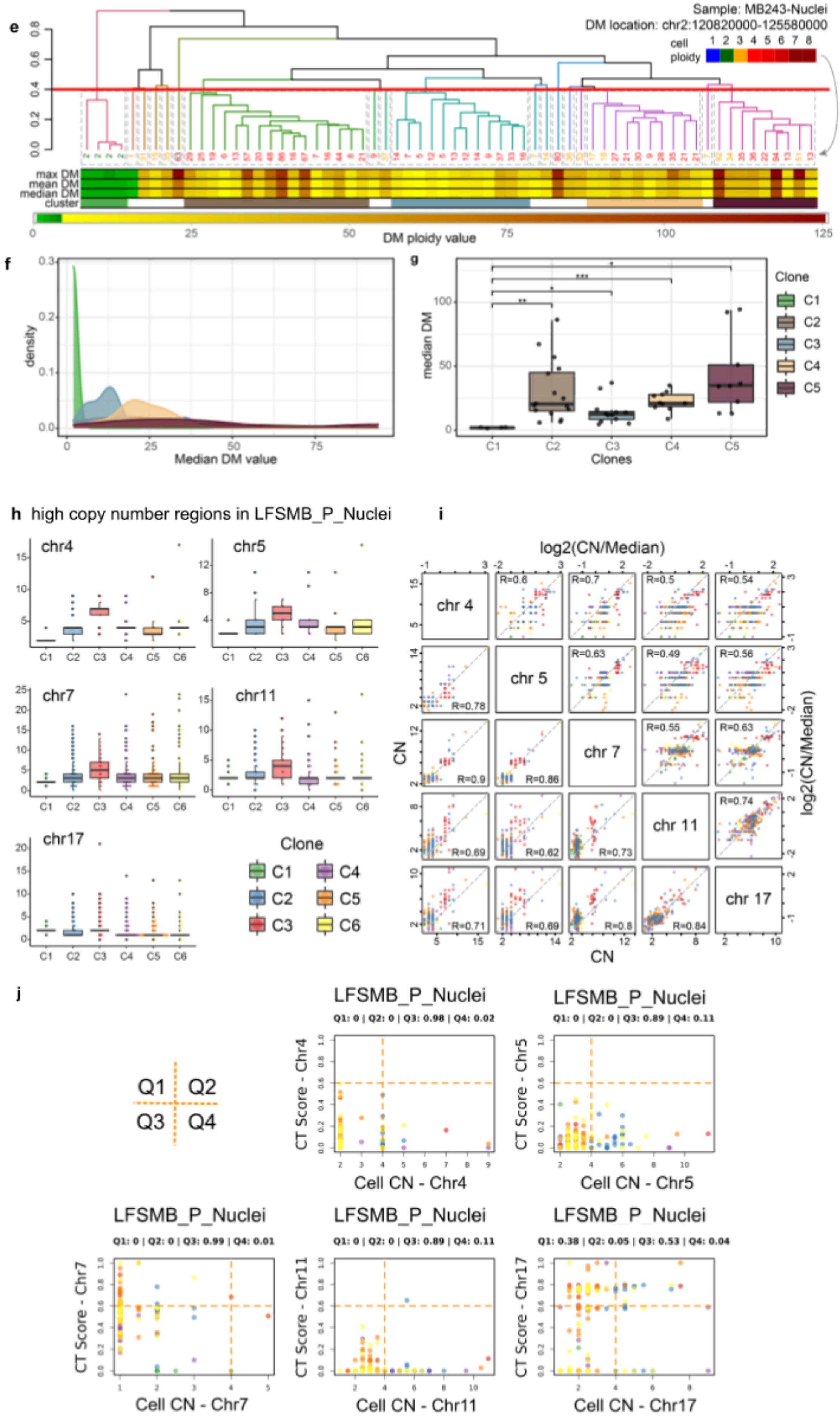

**Supplementary Figure 4. Intra-tumor heterogeneity of druggable genes and double-minute chromosomes. a,** Druggable targets divided into two groups: Druggable genes with focal copy-number gains in all clones except in normal cells and druggable genes with intra-tumor heterogeneity detected by scDNA-seq. In both cases, the log2 ratios between the CNV profile for a given gene locus (as compared to the ploidy for this chromosome) and the median CNV values at the cell level for every clone are shown. Additionally, the ratio between the CNV value at the bulk level and a value of 2 (diploid) is displayed (red triangle). For all genes, expression patterns from the scRNA-seq data are shown for the matched DNA clones. Since several normal cell types were identified in the scRNA-seq data, the reference cell type is sample specific. The sample/reference cell types used are the following: LFS_MBP-Nuclei/Purkinje cells, MB243-Nuclei/astrocytes. Due to the high number of zero values in the nuclei samples, only cells having expression values greater than 0 are contained in the boxplots. Below each boxplot, the number of cells with non-zero expression values in relation to the total is shown. **b,** Heatmaps for chromosomes where potential double-minutes chromosomes are detected in samples RCMB18-PDX and MB243-Nuclei. The double minute chromosome regions are marked with light green rectangles. Validation of circular DNA structures was done in bulk WGS using AmpliconArchitect (see methods). **c,** Copy number distribution of double minute chromosome regions for the different clones in samples RCMB18-PDX and MB243-Nuclei. **d,** Scatter plot comparing the copy number in double-minute regions vs Chromothripsis (CT) Score for the chromosome containing the DM regions across cells in samples RCMB18-PDX and MB243-Nuclei. **e,** Dendrogram after performing hierarchical clustering on cells from the MB243-Nuclei sample. Every leaf in the dendrogram represents a cell, where the label value is the DM ploidy (if multiple bins are affected then the median value is taken) and the color of the label represents the cell ploidy. Below the dendrogram maximum, mean, and median DM ploidy values are shown for each cell, as well as clone assignments. **f,** Density plots of median DM ploidy per clone. **g,** Boxplot of median DM ploidy per clone in MB243-Nuclei, with significance values after performing t-test. Differences for all clone combinations are tested, but only significant p-values are shown. Bonferroni correction is done for multiple-comparison testing. **h,** High copy number regions in LFS_MBP-Nuclei. Putative circular DNA structures were found in bulk WGS using AmpliconArchitect. As compared to other samples, these structures show relatively low copy-numbers (no high amplification as for canonical double minutes regions). **i,** Correlation between CNVs in high CNV regions from different chromosomes in LFS_MBP-Nuclei. Lower triangular matrix contains raw CNV values. Upper triangular matrix contains ploidy corrected CNVs. **j,** Same as panel d but for high CNV regions in different chromosomes in LFS_MBP-Nuclei.

#### Supplementary Figure 5

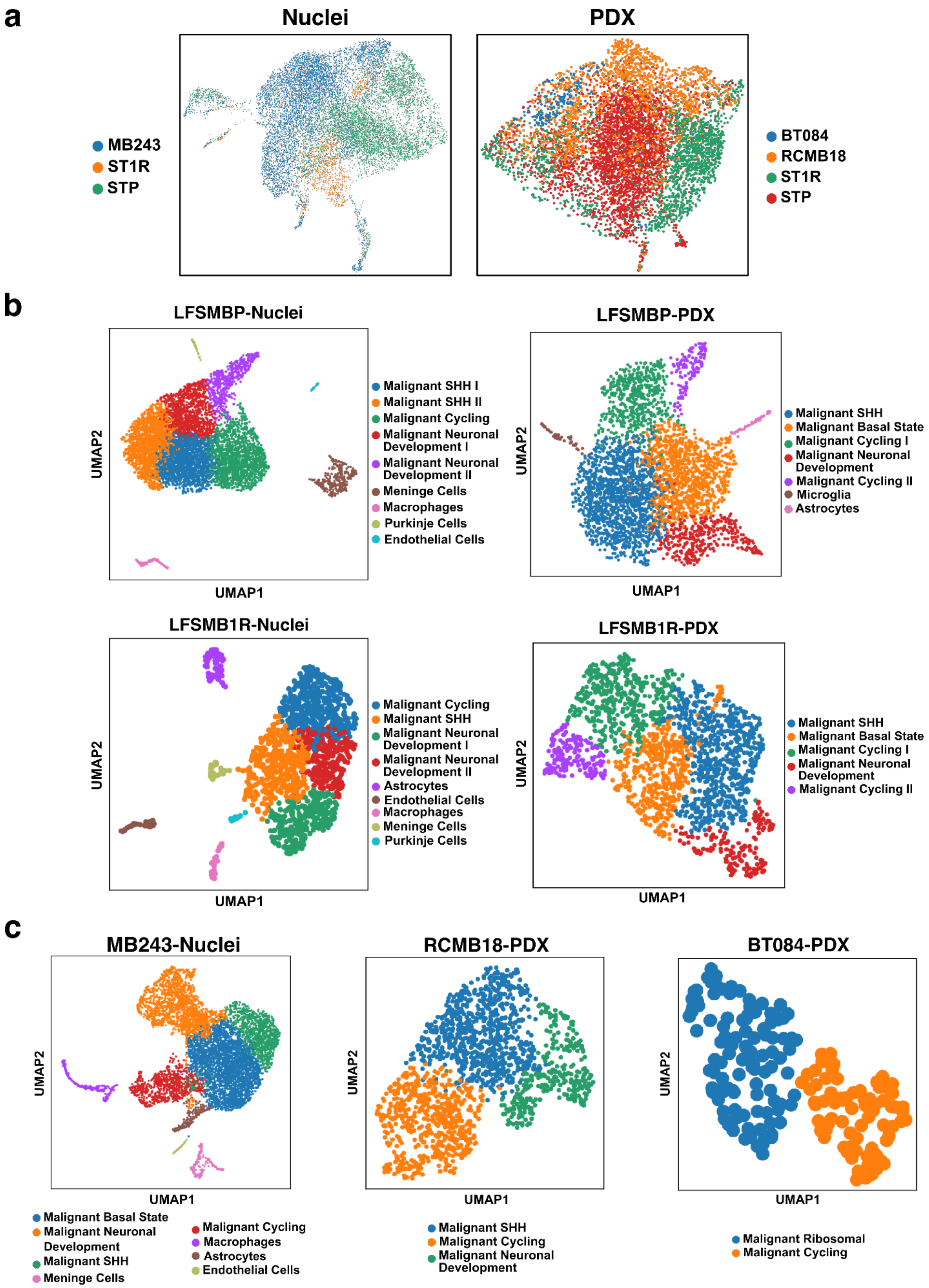

**Supplementary Figure 5. UMAP plots of all analyzed samples depicting cell type identities.** **a,** Aligned UMAP embedding from single-nuclei and single-cell sequencing (PDX) highlighting cells derived from each individual sample. **b,** UMAP embeddings for each individual sample acquired from the main patient of the paper (LFS_MB, including LFS_MBP-Nuclei, 6,669 cells; -PDX, 3,629 cells; and LFS_MB1R-Nuclei, 1,322 cells; -PDX, 1,817 cells). All cells are colored according to their cluster assignment. **c,** UMAP plots for each individual sample acquired from additional patients (MB243-Nuclei, 7,302 cells; RCMB18-PDX, 1,613 cells; BT084-PDX, 182 cells). All cells are colored according to their cell cluster assignment. Clusters were assigned to cell types based on markers from the literature (see Methods).

### Supplementary Figure 6

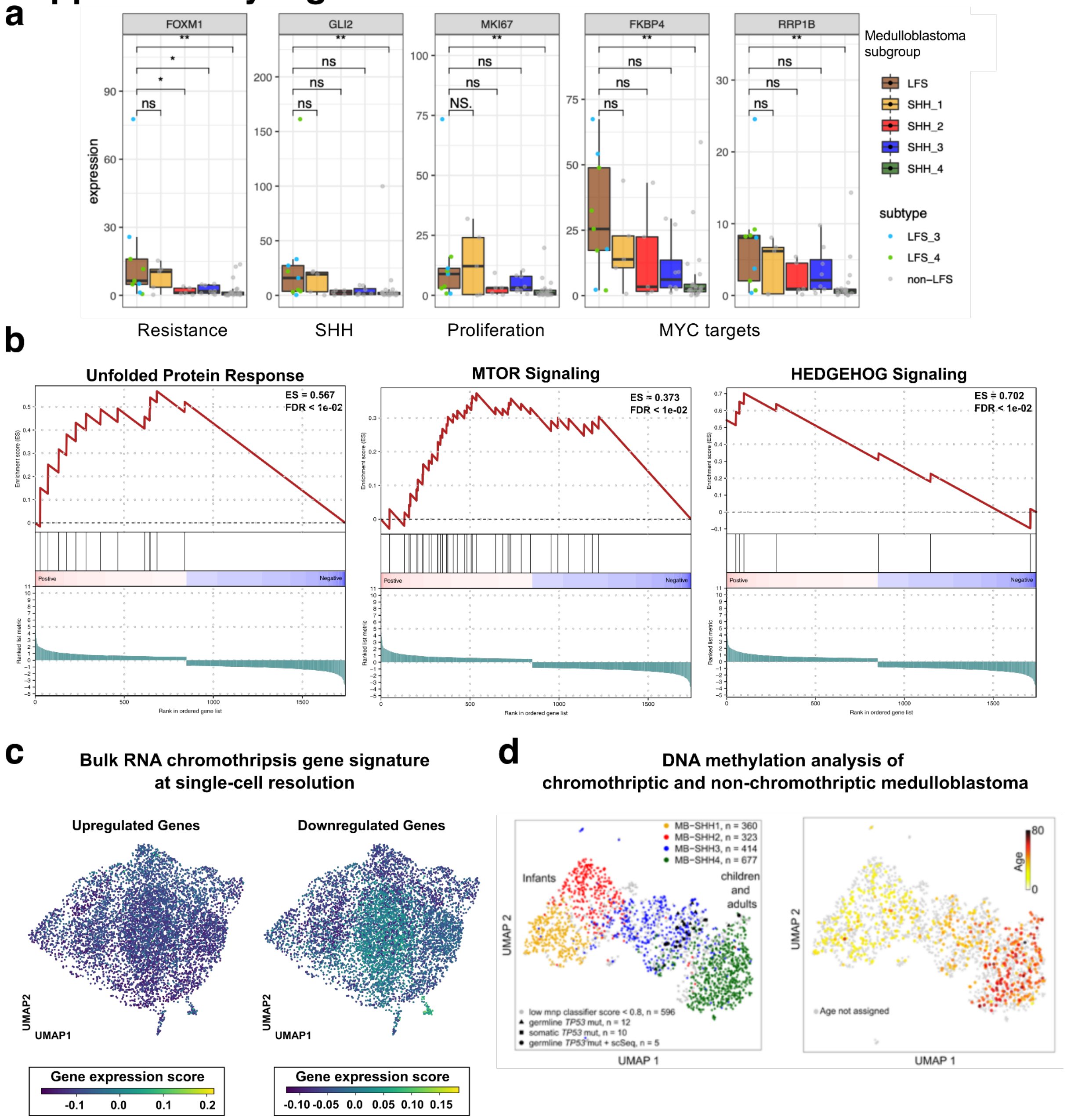


**Supplementary Figure 6. Transcriptional signatures contributing to the aggressiveness of chromothriptic medulloblastomas. a,** Boxplots representing selected genes (*FOXM1*, *GLI2*, *MKI67*, *FKBP4* and *RRP1B*) associated with resistance, SHH, proliferation and *MYC* targets across medulloblastoma subgroups. **b,** Gene Set Enrichment Analysis (GSEA) on differential expression signatures between LFS medulloblastomas versus non-CT SHH from bulk RNA seq data (n=38 non-CT SHH medulloblastomas and 8 LFS medulloblastomas) identified enrichment of Unfolded protein response, *MTOR* signaling and Hedgehog signaling in CT MBs as compared to non-CT MBs (FDR < 0.05, Kolmogorov–Smirnov statistic, Benjamini Hochberg adjusted). **c,** UMAP embedding for the PDX samples profiled with single-cell RNA-seq colored according to the bulk chromothripsis score derived from the differential gene expression analysis shown in Figure **3f**. The genes associated with chromothripsis were split into up- and down-regulated genes. **d,** UMAP embedding results from the analysis of DNA methylation patterns suggesting that CT medulloblastomas developing in LFS patients represent a group between infant and children SHH medulloblastomas, together with SHH MBs showing somatic *TP53* mutations. The plot on the left-hand side is colored according to SHH groups, while the right-hand side shows the age distribution across samples.

### Supplementary Figure 7

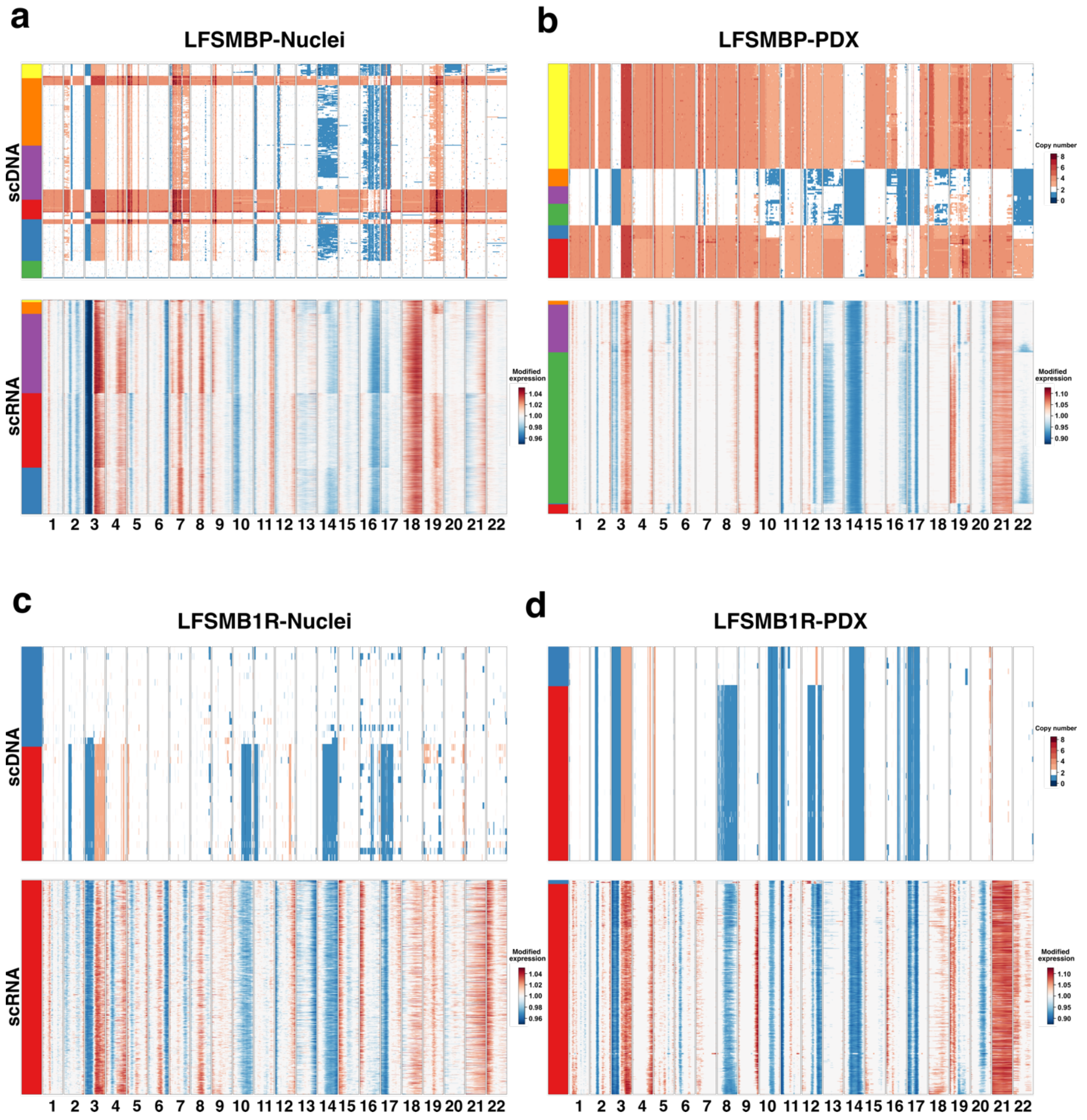

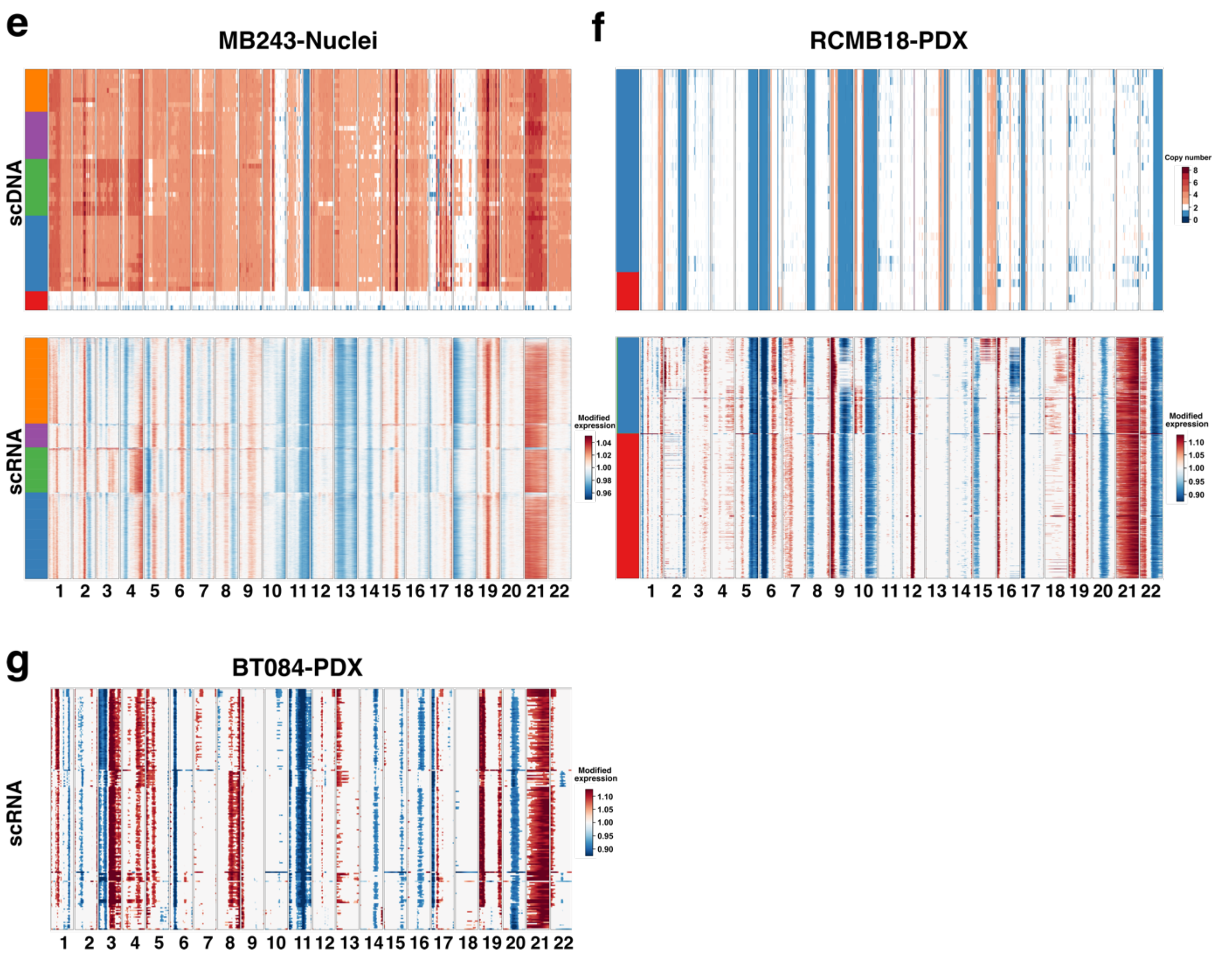


**Supplementary Figure 7. Integrated copy number heatmaps from genomic and transcriptomic data.** Results from scDNA- and scRNA-seq copy number analysis across all samples. The figure shows the scDNA copy number heatmap at the top, while the scRNA-seq copy number heatmap, inferred from inferCNV, is presented at the bottom. In the first part of this figure, samples acquired from the main patient of the paper (LFS_MB, including LFS_MBP-Nuclei, -PDX and LFS_MB1R-Nuclei, -PDX) are shown, while the second part highlights the other four samples (MB243-Nuclei, RCMB18-PDX and BT084-PDX). **a,** Copy number heatmap from scDNA- and scRNA-seq for LFS_MBP-Nuclei. **(Top)** The heatmap is showing the copy number profiles from scDNA. Here, copy number alterations are shown for the 22 autosomal chromosomes, with red representing high copy values, white showing a diploid copy number status and blue depicting low copy number. The scale was cut off at copy number values higher than eight (interval = [0,8]). The color bar at the left-hand side relates to the clones detected in scDNA. Cells were clustered hierarchically within each clone boundary. Importantly, instead of showing copy number values per segment, as shown in **Figure 1**, we here present as transformed to gene-level (see Methods). **(Bottom)** The heatmap highlights the results from the copy number inference based on scRNA-seq data. Copy number profiles were generated using inferCNV, whose output was used to generate an image representation. The color bar at the left-hand side relates to the integrated clones from scDNA, indicating the clone-cell-relationship. Cells were clustered hierarchically within each clone boundary. The scales were limited to [0.95, 1.05] with red representing high modified gene expression, white showing a diploid modified gene expression value of 1 and blue indicating low modified gene expression. For the respective heatmaps, the legends are indicated on the very right hand side. The layout for the subsequent figure panels is represented identical to what is described above. **b, - g,** Results from scDNA- and scRNA-seq copy number analysis for all remaining samples. For BT084-PDX, the analysis from scDNA-seq is missing as this sample was not profiled with this methodology.

### Supplementary Figure 8

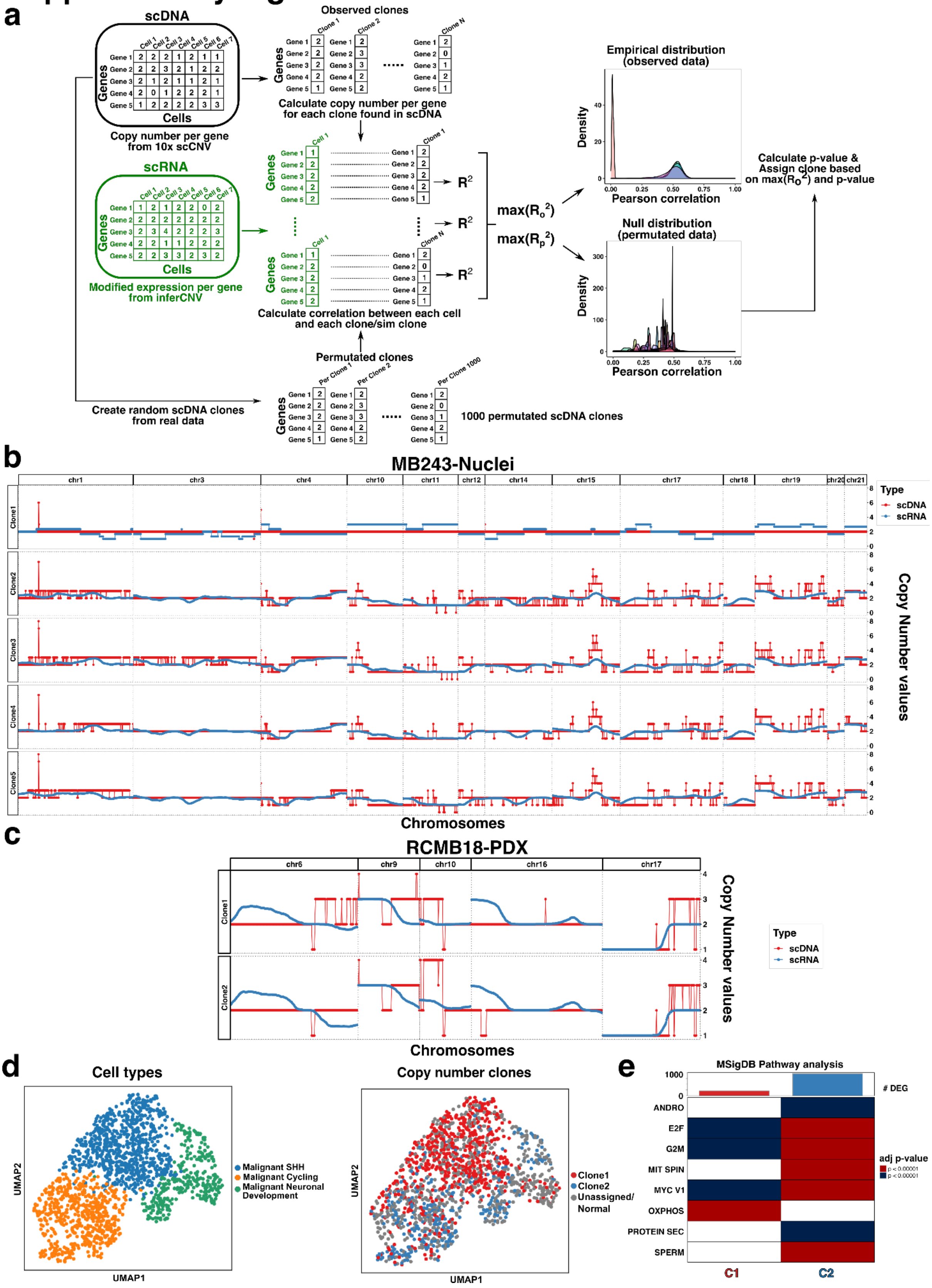

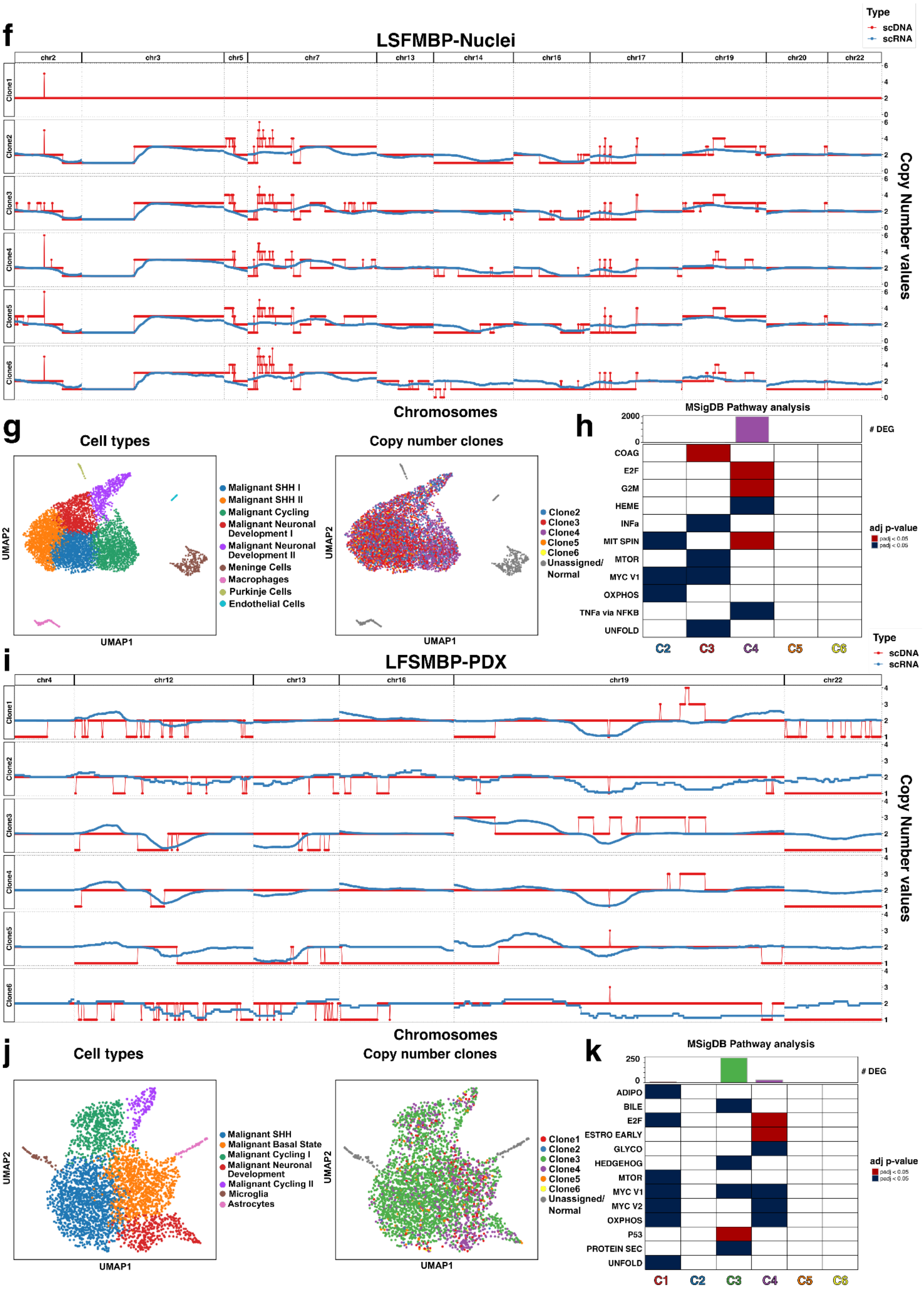
f
LSFMBP-Nuclei
Type
scDNA
scRNA
chr2
chr3
chr5
chr7
chr13
chr14
chr16
chr17
chr19
chr20
chr22
Clone1
Clone2
Clone3
Clone4
Clone5
Clone6
Copy Number values
Chromosomes
g
Cell types
Malignant SHH I
Malignant SHH II
Malignant Cycling
Malignant Neuronal Development I
Malignant Neuronal Development II
Meninge Cells
Macrophages
Purkinje Cells
Endothelial Cells
UMAP1
UMAP2
Copy number clones
Clone2
Clone3
Clone4
Clone5
Clone6
Unassigned/ Normal
h
MSigDB Pathway analysis
2000
0
### DEG
COAG
E2F
G2M
HEME
INFa
MIT SPIN
MTOR
MYC V1
OXPHOS
TNFa via NFKB
UNFOLD
C2
C3
C4
C5
C6
adj p-value
padj < 0.05
padj < 0.05
i
LFSMBP-PDX
Type
scDNA
scRNA
chr4
chr12
chr13
chr16
chr19
chr22
Clone1
Clone2
Clone3
Clone4
Clone5
Clone6
Copy Number values
Chromosomes
j
Cell types
Malignant SHH
Malignant Basal State
Malignant Cycling I
Malignant Neuronal Development
Malignant Cycling II
Microglia
Astrocytes
UMAP1
UMAP2
Copy number clones
Clone1
Clone2
Clone3
Clone4
Clone5
Clone6
Unassigned/ Normal
k
MSigDB Pathway analysis
250
0
### DEG
ADIPO
BILE
E2F
ESTRO EARLY
GLYCO
HEDGEHOG
MTOR
MYC V1
MYC V2
OXPHOS
P53
PROTEIN SEC
UNFOLD
C1
C2
C3
C4
C5
C6
adj p-value
padj < 0.05
padj < 0.05

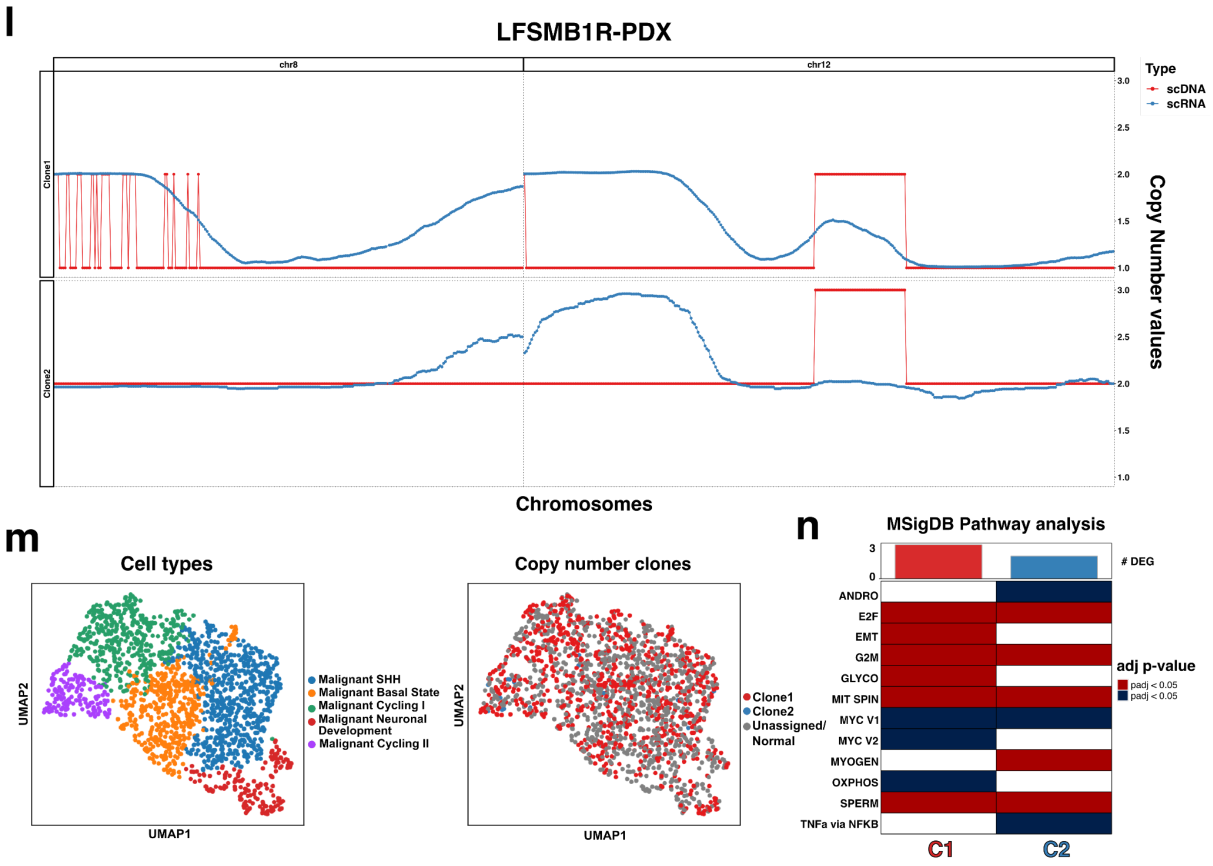


**Supplementary Figure 8. Downstream comparison and consequences of the copy number integration of scRNA- and scDNA-seq data.** Results from the copy number integration and downstream transcriptomic analysis for all samples with matched scRNA- and scDNA-seq data. **a,** Technical schematic of the computational strategy for aligning scRNA-seq profiles to CNV profiles of individual clones. **b,** Integration results for MB243-Nuclei from the procedure described in **Figure 7a**. In this visualization, the variable chromosomes selected for the integration are shown on the x-axis, while the copy number values per clone from scDNA-seq are shown on the y-axis. Highlighted are the aggregated pseudo bulk profiles from scDNA (red) and scRNA (blue). The copy number values from the scDNA data were normalized to a median diploid copy number. **c,** Integration results for RCMB18-PDX, equivalent to the plot shown in **a** panel. **d,** UMAP embedding depicting 1,613 cells from scRNA-seq after quality checks. The plot shows the individual cells colored according to the cell type identity as well as according to the clones from the integrated analysis. Phenotypically normal cells as well as cells which did exceed a p-value > 0.05 were excluded and are marked as unassigned or normal. **e,** Heatmap for pathway enrichment analysis for each individual clone detected in scRNA-seq. Additionally, the number of significantly differentially expressed genes comparing the cells of a specific clone with all other cells, is shown as a bar plot on the top. Genes are considered significantly differentially expressed if they have a p-value < 0.05 (FDR < 0.05, Wilcoxon rank sum test, Benjamini Hochberg adjusted, one clone versus all). Each bar is colored based on the clone color shown in **b**. For the pathways, significance is shown as p-value < 0.05 (FDR < 0.05, Kolmogorov–Smirnov statistic, Benjamini Hochberg adjusted). Depending on the type of differential expression (up- or down-regulation), the pathways are shown in shades of red or blue. **f, - h,** Integration results for LFS_MBP-Nuclei, with the schematic following the identical description given in **b - e**. **i, - k,** Integration results for LFS_MBP-PDX. **l, - n,** Integration results for LFS_MB1R-PDX. Abbreviations - ADIPO: Adipogenesis, ANDRO: Androgen response, BILE: Bile acid metabolism, COAG: Coagulation, G2M: G2M checkpoint, GLYCO: Glycolysis, HEME: Heme metabolism, HEDGEHOG: Sonic hedgehog signaling, PROTEIN SEC: Protein secretion, OXPHOS: Oxidative phosphorylation pathway, SPERM: Spermatogenesis, ESTRO EARLY: Estrogen response early, INFa: Interferon-a response, MYOGEN: Myogenesis, E2F:

E2F targets, MTOR: MTOR signaling, MYC V1 & MYC V2: MYC signaling, P53: P53 pathway, EMT: Epithelial mesenchymal transition, TNFa via NFKB: TNFa via NFKB signaling, UNFOLD: Unfolded protein response

### Supplementary Figure 9

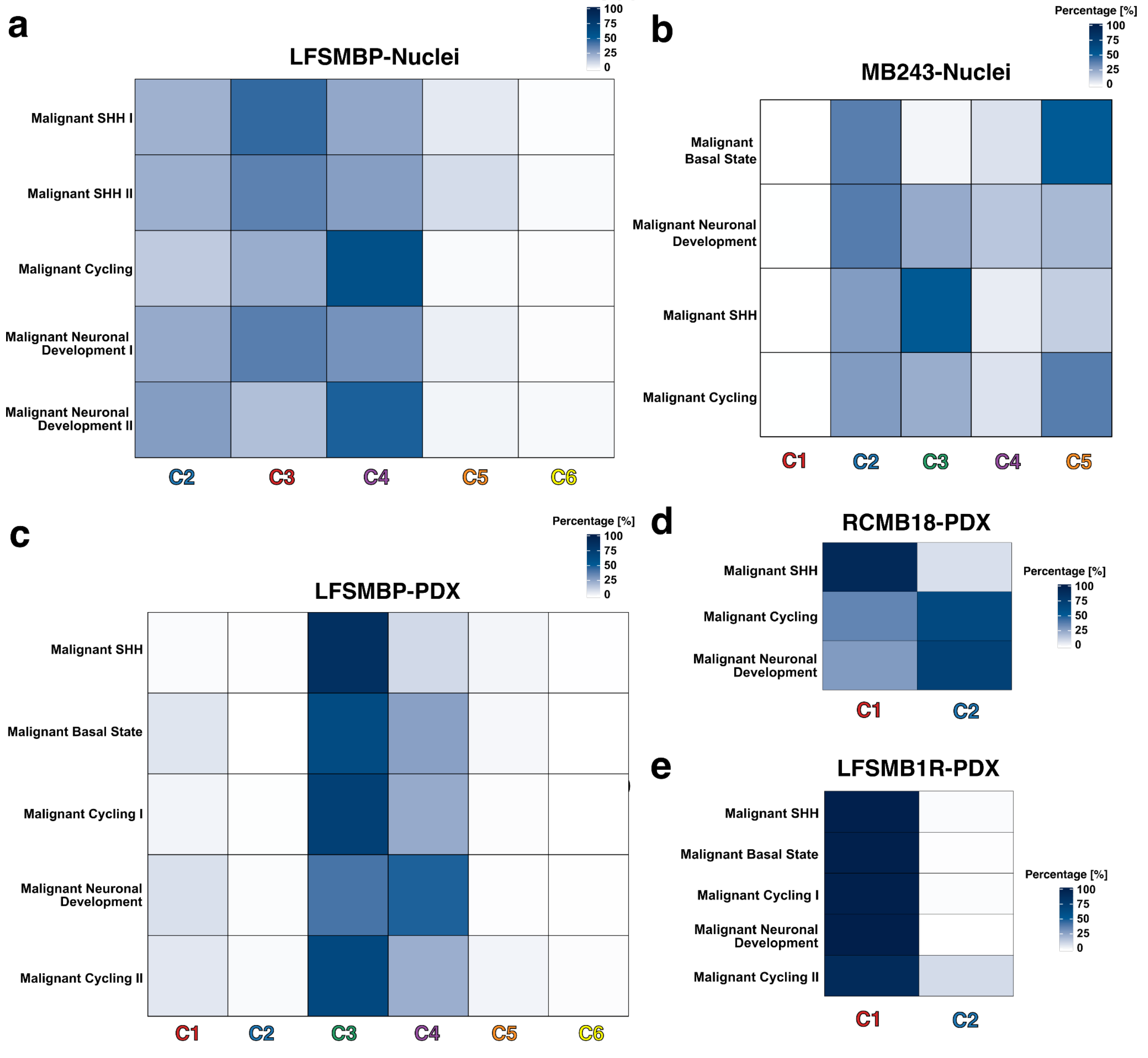


**Supplementary Figure 9. Association between copy number clones and cell type identity.** Visualized contingency tables for intersection between integrated copy number clones and cell type assignments for all samples with more than one clone. **a,** Contingency table for LFS_MBP-Nuclei highlighting the proportion of the cell type population, which has been assigned to a specific copy number clone. Only malignant cell types, for which the copy number profile could be determined, are shown on the y-axis, while the copy number clones are present on the x-axis. The percentage of cells for each clone is highlighted in blue, with white representing 0% and dark blue 100%. **b,** Contingency table equivalent to **a** for MB243-Nuclei. **c,** Contingency table equivalent to **a** for LFS_MBP-PDX. **d,** Contingency table equivalent to **a** for RCMB18-PDX. **e,** Contingency table equivalent to **a** for LFS_MB1R-PDX.

#### Supplementary Figure 10

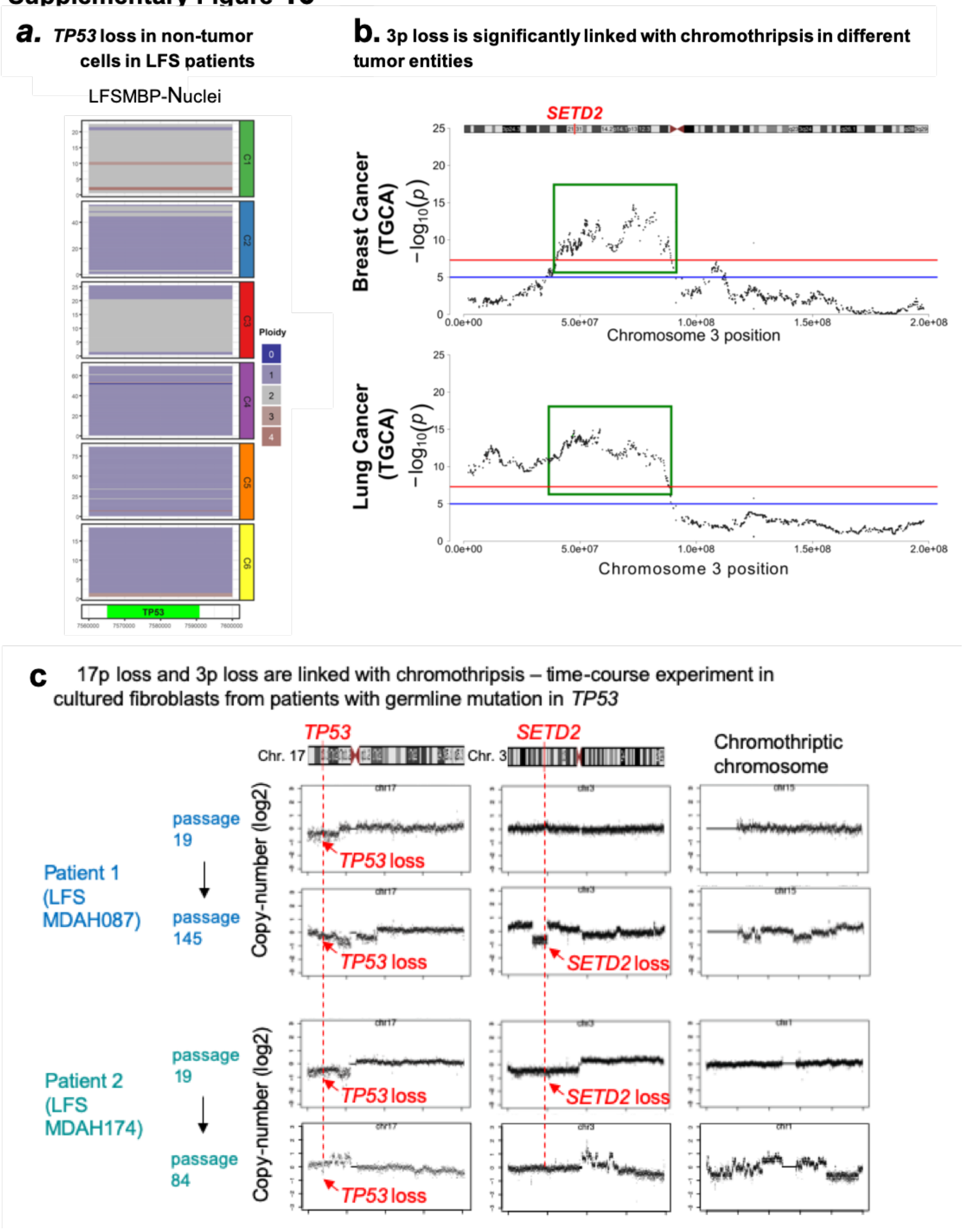


**Supplementary Figure 10. Loss of chr3p and chr17p as early events linked to chromothripsis. a,** *TP53* loss in rare non-tumor cells in LFS patients. **b,** Chromosome 3p loss is significantly linked with chromothripsis in different tumor entities. **c,** 17p and 3p loss are linked with chromothripsis - time course experiment in cultured fibroblasts from patients with germline mutation in *TP53*.

#### Supplementary Tables

**Supplementary Table 1. Overview of quality control and filtering criteria in scDNA-seq.**

**Supplementary Table 2. Chromothripsis scoring for all samples** (chromothripsis scoring method for clones by majority vote CNV profiles aggregation as explained in the methods section).

**Supplementary Table 3. Overview of quality control and filtering criteria in scRNA-seq as well as integration details.**

**Supplementary Table 4. Results of the bulk RNA-seq differential gene expression analysis using DESeq2.**
